# Divergent and diversified proteome content across a serially acquired plastid lineage

**DOI:** 10.1101/2022.11.30.518497

**Authors:** Anna M. G. Novák Vanclová, Charlotte Nef, Adél Vancl, Fuhai Liu, Zoltán Füssy, Chris Bowler, Richard G. Dorrell

## Abstract

Dinoflagellates are a diverse group of ecologically significant micro-eukaryotes, prone to the loss and replacement of their plastids via serial endosymbiosis. One such replacement took place in the ancestors of the Kareniaceae, which harbor haptophyte-derived plastids containing the pigment fucoxanthin, instead of the ancestral peridinin dinoflagellate plastid. The metabolic functions of the fucoxanthin plastid are performed by a diverse range of nucleus-encoded and plastid-targeted proteins, which may originate from the haptophyte ancestor of the fucoxanthin plastid, the peridinin-containing ancestor of the dinoflagellate host, and/or from lateral gene transfers. However, the evolutionary composition of fucoxanthin plastid proteomes across the diversity of kareniacean dinoflagellates remains poorly understood. Here, we determine the total composition of the plastid proteomes of seven distantly-related kareniacean dinoflagellates, including newly-sequenced members of three genera (*Karenia, Karlodinium*, and *Takayama*). Using a custom plastid-targeting predictor, automatic single-gene tree building and phylogenetic sorting of plastid-targeted proteins, we project relatively limited (~10%) and functionally distinctive contributions of the haptophyte endosymbiont to the fucoxanthin plastid proteome, in comparison to plastid-targeted proteins of dinoflagellate host origin. Considering a concatenated multigene phylogeny of haptophyte-derived plastid proteins, we show that the haptophyte order Chrysochromulinales is the closest living relative of the fucoxanthin plastid donor. We additionally perform detailed analyses of the N-terminal targeting sequences of kareniacean plastid signal peptides, reporting a surprisingly high sequence conservation. Finally, considering planetary-scale distributions of key kareniacean genera and haptophyte orders from *Tara* Oceans, we suggest ecological and mechanistic factors accompanying the endosymbiotic acquisition of the fucoxanthin plastid.

## Introduction

The origin of oxygenic photosynthesis, and its adoption by eukaryotes, has fundamentally changed the evolutionary landscape of life. Originating in cyanobacteria, photosynthesis has spread across the eukaryotes through multiple endosymbiotic acquisitions of plastids (or chloroplasts). These began with the primary plastids of cyanobacterial origin that represent the defining synapomorphy of Archaeplastida (glaucophytes, red and green algae, and plants) and are surrounded by two membranes. Plastids subsequently spread into other groups (e.g., diatoms, haptophytes, dinoflagellates, euglenids) through multiple endosymbiotic acquisitions of eukaryotic algae, forming “complex plastids” that possess additional membranes, often contained within the endoplasmic reticulum. Across the tree of life, plastids provide their hosts with novel metabolic and biosynthetic capabilities and serve as important vectors of horizontal gene transfer between different algal lineages (Howe et al., 2008; Ku et al., 2015; Novák Vanclová et al., 2019; Dorrell et al., 2021).

Plastids have their own genomes that typically encode core photosynthetic proteins alongside some of their genetic housekeeping machinery and primary metabolic enzymes. Their size and coding capacity vary, typically possessing 20-250 genes, driven by gradual and somewhat convergent gene loss (Uthanumallian et al., 2021). Plastids however require a much larger inventory of proteins to function (e.g., > 2000 in *Arabidopsis* and model eukaryotic algal groups; Kleffmann et al., 2004; Gruber et al., 2015; Novák Vanclová et al., 2019); and most of the individual plastidial proteins are encoded in the nucleus and post-translationally imported based on a cleavable N-terminal targeting signal. These plastid-targeted proteins comprise the expression products of plastid-derived genes that were transferred to host nucleus, host-derived genes newly adapted for plastidial function, and genes originating from lateral gene transfer (LGT) from other sources, including other symbionts that the host has interacted with over its evolutionary history (Moustafa et al., 2008). This makes all plastid proteomes innately chimeric entities, or “shopping bags”, whose composition may reflect both general principles of organelle evolution and the unique evolutionary histories of individual lineages (Larkum et al., 2007).

Dinoflagellates are a diverse group of unicellular algae that are ecologically and economically significant; including crucial symbionts of corals, and toxin-producing species that can cause major losses in fisheries and endanger public health (Hackett et al., 2004; Wang, 2008). Within the eukaryotic tree of life, they fall among the supergroup Alveolata, specifically Myzozoa along with chrompodellids and apicomplexans (Ševčíková et al., 2015). Dinoflagellates are of further evolutionary interest given their unique and idiosyncratic cell biology. Their nuclear genomes are generally extremely large (up to several hundred Gbp) with permanently condensed chromosomes that lack histones, while their mitochondrial genomes are very streamlined (Lin, 2011; Dorrell and Howe, 2015). The evolutionarily ancestral dinoflagellate plastids are descended from red algae and bound by three membranes (Moog and Maier, 2017). The ancestral dinoflagellate plastid is further marked by multiple functional oddities, including a unique carotenoid pigment, peridinin (Haxo et al., 1976); and a bacterial-like nucleus-encoded form II RuBisCO (Morse et al., 1995) in contrast to the partially plastid-encoded form I RuBisCO found in all other plastid lineages. Dinoflagellate plastid genomes are highly fragmented and organized into small unigenic elements termed “minicircles”. These transcripts undergo distinctive maturation events, including extensive substitutional editing and the addition of a 3’ poly(U) tail not found in most other plastid lineages (Zauner et al., 2004; Wang and Morse, 2006; Dorrell and Howe, 2015).

Dinoflagellates as a group are ecologically opportunistic: most plastid-bearing representatives are in fact mixotrophic (Cohen et al., 2021; Jeong et al., 2021) and in global meta-genomic analyses (e.g., *Tara* Oceans) dinoflagellates appear mostly as predators and parasites (Pierella Karlusich et al., 2022). Their plastids are frequently secondarily reduced to non-photosynthetic organelles, lost completely (Saldarriaga et al., 2001; Sanchez-Puerta et al., 2007; Gornik et al., 2015; Cooney et al., 2022), or replaced in new symbiogenetic events. These may result in fully integrated plastids (Tengs et al., 2000; Matsumoto et al., 2011), complex endosymbionts retaining plastids, mitochondria, and nuclei (Imanian and Keeling, 2007; Sarai et al., 2020), and kleptoplastids of various degrees of stability (Koike et al., 2005; Myung et al., 2006; Gast et al., 2007). In some cases, the co-existence of a new organelle or endosymbiont with a remnant of the ancestral plastid has been proposed (Hehenberger et al., 2019), and in one lineage, a complete repurposing of parts of the plastid and mitochondrion into an enigmatic, photosynthesis-unrelated optical structure termed “ocelloid” took place (Gavelis et al., 2015).

Kareniaceae, typified by the genera *Karenia, Karlodinium*, and *Takayama*, are mixotrophic dinoflagellates whose plastids are related to those of haptophytes. Kareniaceae are the most abundant dinoflagellate lineage with serially acquired plastids in *Tara* Oceans (Table S1, based on data from Engelen et al., 2015), and both *Karenia* and *Karlodinium* have important impacts as toxic components of harmful algal blooms. Further ecological and evolutionary complexity found within the Kareniaceae is the Antarctic native Ross Sea dinoflagellate (RSD) (Gast et al., 2007; Hehenberger et al., 2019), which uses a kleptoplastid from the haptophyte *Phaeocystis antarctica* acquired independently to that of other Kareniaceae.

Instead of peridinin, plastids of the Kareniaceae contain the accessory pigment fucoxanthin typical of their haptophyte ancestors (Zapata et al., 2004, 2012), and are bound by four membranes, the outermost of which is continuous with the endoplasmic reticulum as per haptophytes, and distinct from the three membrane-bound plastids found in peridinin dinoflagellates.

Partial kareniacean plastid genomes and transcriptomes have been sequenced for the species *Karlodinium micrum* (syn. *veneficum*) and *Karenia mikimotoi*, indicating the retention of around 100 genes on a single circular or linear chromosome (Gabrielsen et al., 2011; Dorrell et al., 2016), which is somewhat fewer than the ca. 140 genes associated with haptophyte plastids, alongside the presence of episomal minicircles (Richardson et al., 2014). Further studies, based on expressed sequence tags (ESTs) (Ishida and Green, 2002; Nosenko et al., 2006; Patron et al., 2006) and, more recently, transcriptomes of *Karenia brevis* and *Karlodinium micrum* realized through the Marine Microbial Eukaryote Transcriptome Sequencing Project (MMETSP) (Burki et al., 2014; Keeling et al., 2014; Matsuo and Inagaki, 2018; Hehenberger et al., 2019), have provided foundational insights into the nucleus-encoded proteome of the kareniacean plastid. These are defined by a bipartite targeting sequence, consisting of an N-terminal signal peptide followed by a hydrophilic transit peptide, similar to those of haptophytes, although with apparent differences in composition (Patron and Waller, 2007; Yokoyama et al., 2011). Phylogenetic analysis of the genes encoding plastid-targeted kareniacean proteins (Nosenko et al., 2006; Patron et al., 2006; Burki et al., 2014; Bentlage et al., 2016; Matsuo and Inagaki, 2018) indicate that some resolve with haptophytes (i.e., the endosymbiont ancestor), while others show alternative origins. These include key elements of the plastid gene expression machinery (e.g., RNA editing and poly(U) tail addition), which are likely to have been derived from the dinoflagellate host and potentially associated with the ancestral peridinin plastid (Dorrell and Howe, 2012; Jackson et al., 2013); and cofactor (e.g., isopentenyl pyrophosphate and protoporphyrin/haem) biosynthetic pathways, which resolve principally either with dinoflagellates or sources suggesting horizontal acquisition (Matsuo and Inagaki, 2018).

To date, a reconstruction of the entire fucoxanthin plastid has yet to be attempted, and its overall metabolic composition and specific evolutionary origin within the haptophytes remain unresolved (Yoon et al., 2002; Takahashi et al., 2019; Leblond et al., 2022). Here, we present a holistic reconstruction of the pan-kareniacean plastid proteome across seven model species, integrating newly sequenced transcriptomic datasets for two additional strains of *Karenia* and *Karlodinium* and the first transcriptome for the genus *Takayama*, *in silico* identification and phylogenomic analysis of genes encoding plastid-targeted proteins.

These data elucidate the metabolic functions of the fucoxanthin plastid in its kareniacean host, its deep phylogenetic origins, and the evolutionary principles that may have constrained its genesis and early history. In particular, we emphasize the significance of integration with pre-existing components and systems inherited from the host and its previous plastid. Our study illuminates the deep evolutionary history of this ecologically important algal lineage, and some of the functional principles that may underpin symbiogenetic plastid establishment across eukaryotes.

## Results

### Pan-transcriptomics reveals the core fucoxanthin plastid proteome

Five new transcriptomic datasets were produced for fucoxanthin dinoflagellates: *Karenia mikimotoi* and *Karenia papilionacea*, the RCC3446 strain of *Karlodinium micrum* and its distant relative *Karlodinium armiger*, and for *Takayama helix*, a predator of other dinoflagellates more closely related to *Karlodinium* than *Karenia* (Jeong et al., 2016), for which to our knowledge this is the first reported transcriptomic dataset. These libraries, alongside four previously sequenced MMETSP libraries for *Karenia brevis*, one for *Karlodinium micrum* strain CCMP2283, and an independently sequenced transcriptome for RSD, represent the pan-kareniacean library used for all downstream analyses.

The raw dataset sizes range from around 200 to 660 Mbp but contain large amounts of redundant or partially redundant transcripts, reflecting the extensive paralogization and pseudogenization associated with dinoflagellate nuclear genomes (Stern et al., 2010). To reduce the size and redundancy of our datasets for subsequent analysis, we selected the most complete paralogs of each studied protein by cd-hit. The completeness of the datasets as assessed by BUSCO is around 70% which is rather high in comparison to other dinoflagellate datasets (Table 1, Table S2).

**Table 1:**
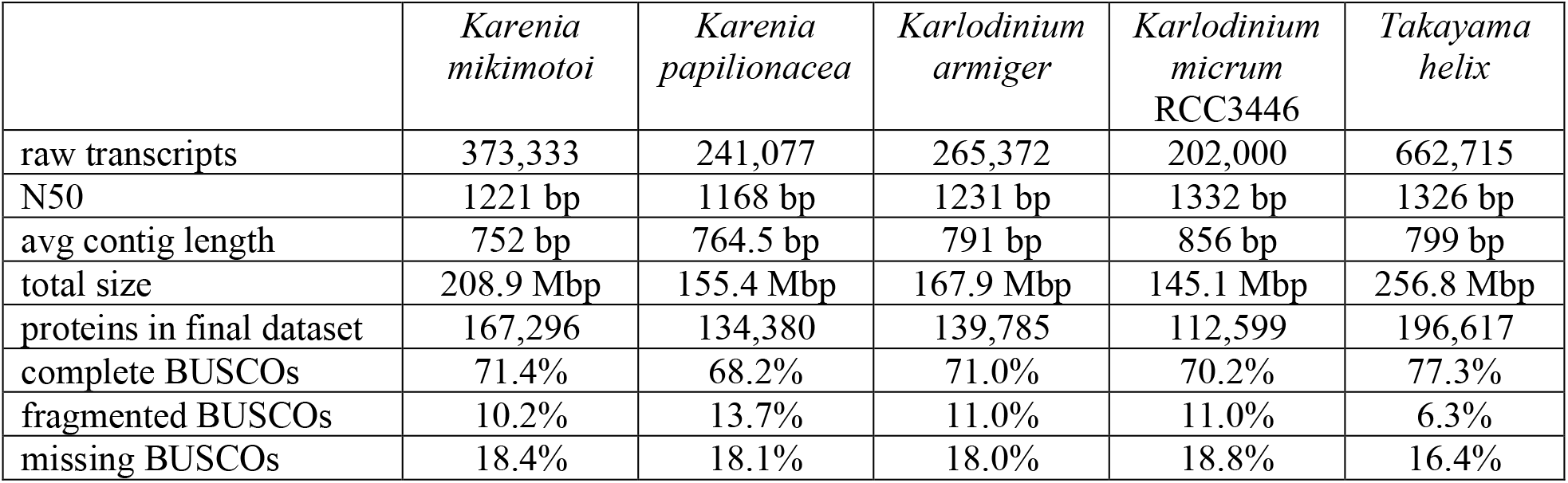
Assembly statistics and assessment of completeness for the five new transcriptomes.

|  | *Karenia mikimotoi* | *Karenia papilionacea* | *Karlodinium armiger* | *Karlodinium micrum* RCC3446 | *Takayama helix* |
| --- | --- | --- | --- | --- | --- |
| raw transcripts | 373,333 | 241,077 | 265,372 | 202,000 | 662,715 |
| N50 | 1221 bp | 1168 bp | 1231 bp | 1332 bp | 1326 bp |
| avg contig length | 752 bp | 764.5 bp | 791 bp | 856 bp | 799 bp |
| total size | 208.9 Mbp | 155.4 Mbp | 167.9 Mbp | 145.1 Mbp | 256.8 Mbp |
| proteins in final dataset | 167,296 | 134,380 | 139,785 | 112,599 | 196,617 |
| complete BUSCOs | 71.4% | 68.2% | 71.0% | 70.2% | 77.3% |
| fragmented BUSCOs | 10.2% | 13.7% | 11.0% | 11.0% | 6.3% |
| missing BUSCOs | 18.4% | 18.1% | 18.0% | 18.8% | 16.4% |

### Guided *in silico* prediction reveals within-lineage divergence in fucoxanthin plastid proteome contents

A modified version of ASAFind (Gruber et al., 2015), which has previously been used to reconstruct *in silico* the haptophyte plastid proteome (Dorrell et al., 2017), was constructed with a custom scoring matrix for fucoxanthin dinoflagellate plastids (see Materials and Methods) and used in combination with SignalP 5.0 in the final prediction pipeline. After preliminary automatic annotation of the retrieved *in silico* plastidial proteomes of all studied organisms by KAAS (https://www.genome.jp/kegg/kaas/; Moriya et al., 2007), we noticed that some of the integral plastidial proteins missing from these datasets could be captured by an alternative prediction approach combining PrediSI (Hiller et al., 2004) and ChloroP (Emanuelsson et al., 1999) employed previously for *Euglena gracilis* data (Ebenezer et al., 2017), and so included these as well. The proportion of redundancy-treated translated transcripts predicted as plastid-targeted ranged from 7.5 to 14.5 % in *K. micrum* and *K.brevis*, respectively (~9.5 % being both average and modal value). Of these, most have a targeting sequence predictable by modified ASAFind, while between 16 to 22% represent additions based solely on the abovementioned alternative signal.

The putative phylogenetic origin of all predicted plastid proteins was investigated via a custom pipeline, integrating homology mining, single-gene tree construction and sorting based on topology and its statistical support (Figure S1). Between 40 and 55 % of each proteome dataset had some homologs in our reference database containing representatives for most major eukaryotic groups, particularly enriched in haptophyte and dinoflagellate transcriptomes (see Materials and Methods). In about one third of these, the identified homologs were shared solely with other dinoflagellates and were therefore considered dinoflagellate-specific proteins without any further phylogenetic analysis. For the rest, single-gene trees (29,329 in total) were constructed and automatically sorted based on which evolutionary origin they support. The main two investigated categories were “plastid-early” (i.e., vertically inherited from the dinoflagellate host; colour-coded as blue and purple dependent on whether the protein retrieved only dinoflagellate homologues or showed deeper homology to other alveolates or myzozoans lineages) and “plastid-late” (i.e., immediately clustering with the haptophyte endosymbiont; colour-coded as orange in all graphical materials). In addition to this, potential cases of LGT from prokaryotes, green algae and ochrophytes were identified.

Of all the proteins with homologs in the database, 65-80% were categorized as “plastid-early” in all studied organisms (Figure 1A). The proportion of proteins categorized as “plastid-late” is generally in accord with the previously published estimates (Burki et al., 2014) but there was a noticeable difference between the genera, with approximately 15% noted in all three species of *Karenia*, 10% in *Karlodinium*, and 6% in *Takayama*. The proportion of proteins shared and clustering with ciliates, i.e., primarily heterotrophic alveolates (“non-plastid signal”), is almost negligible in all studied organisms and so is the contribution of prokaryotic or “green” LGT (cases of which do not exceed 20 per organism). The amount of “brown” LGT is slightly higher than the amount of non-plastid signal (varying from 12 to 107 per organism). Between 10 and 20 % of signal in each organism was not sortable into any of the investigated categories.

**Figure 1:**
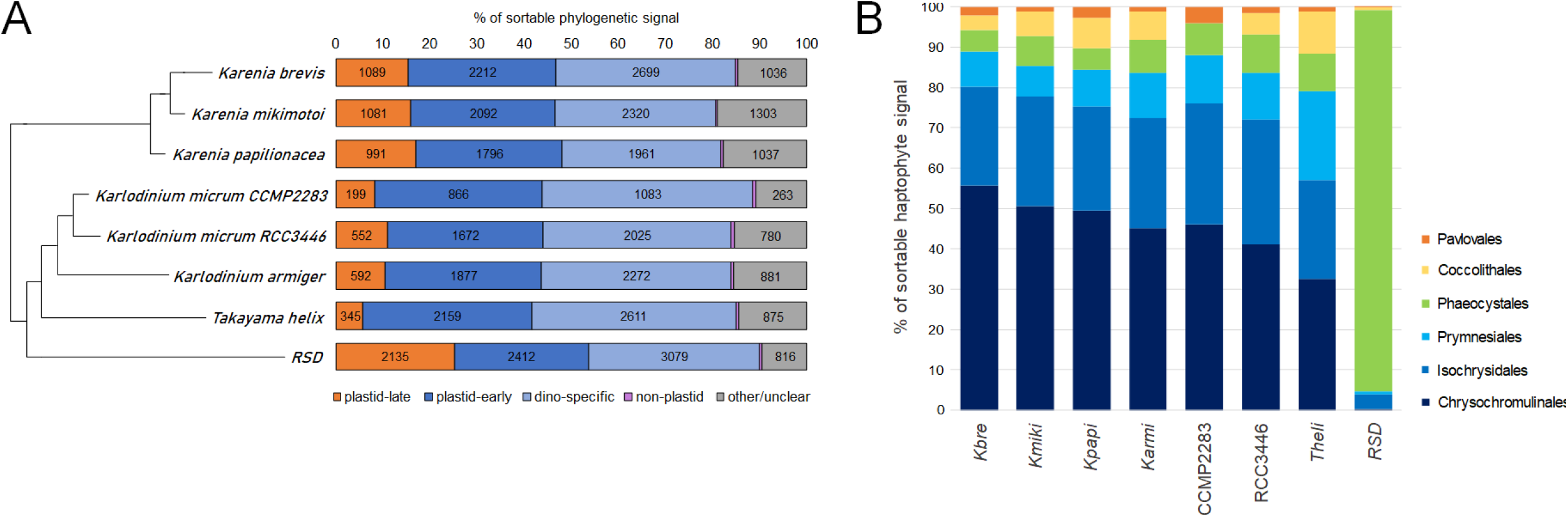
Ratios of phylogenetic signal of plastid proteins with detectable homologs: plastid-late (haptophyte-like; orange), plastid-early (dinoflagellate-like or dinoflagellate-specific; blue), ancestrally alveolate-like (purple), and other or unclear (grey) (A), and further breakdown of the plastid-late signal traceable to a concrete haptophyte family (Chrysochromulinales (dark blue), Isochrysidales (blue), Prymnesiales (light blue), Phaeocystales (green), Coccolithales (yellow), and Pavlovales (orange)) (B).

### Phylogenomics reveals a chrysochromulinacean and potentially incongruent origin of the fucoxanthin plastid

Previous studies of the fast-evolving fucoxanthin plastid genome revealed a deep-branching position sister to all prymnesiophytes but no affinity towards individual orders (Choi et al., 2017; Klinger et al., 2018; Kawachi et al., 2021). To recover a more specific affinity, we enumerated the plastid-targeted proteins clustering specifically with one of six haptophyte families (Chrysochromulinales, Isochrysidales, Phaeocystales, Coccolithales, Pavlovales, and Prymnesiales; Figure 1B), constituting approximately one-quarter of the total plastid-late signal.

Across all tree topologies, we note of selected tree topologies revealed frequent apparent non-monophyly of the kareniacean genes, with *Takayama* sequences often branching separately from the others. Nonetheless, for each of the seven studied fucoxanthin-containing Kareniaceans the most abundant category (up to >50%) was the Chrysochromulinales. A concatenation of 22 genes that were universally identified as plastid-late across all fucoxanthin genera, independent of deeper haptophyte origin, and were either present in a single copy or had only a few easily identifiable and removable paralogs, further robustly placed the fucoxanthin plastid as a sister-group to the Chrysochromulinales (sp. *Chrysochromulina brevifilum*, *Chrysochromulina rotalis* and *Chrysochromulina sp*.; 100% bootstrap support), to the exclusion of the closest related haptophyte taxa within the Prymnesiales (Figure 2A).

**Figure 2:**
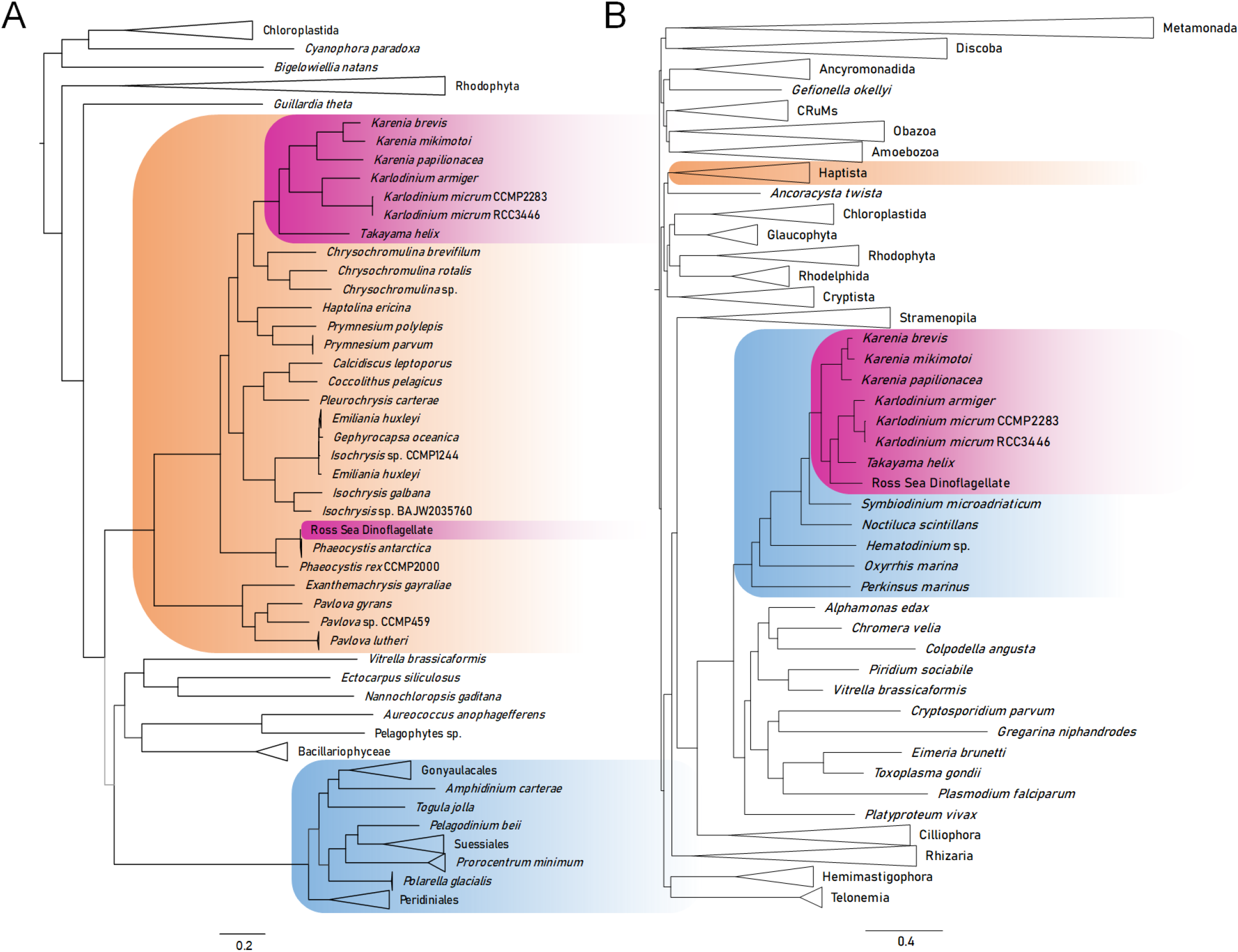
The eight kareniaceans in the pan-eukaryotic phylogenetic context as reconstructed by IQ-TREE based on matrix of 241 genes prepared using PhyloFisher toolkit (A) versus the plastidial phylogeny reconstructed based on 22 proteins of previously determined plastid-late origin (B); bootstrap support is expressed by the branch colour (black for ≥90%, dark grey for ≥75%, light grey for <75%). While RSD places sister to its kleptoplastid donor *Phaeocystis antarctica*, the three non-RSD kareniaceans form a monophyletic clade sister to Chrysochromulinales with maximum bootstrap support on all branches. Inside this clade, *Karenia* and *Karlodinium* cluster together to the exclusion of *Takayama*.

The plastid-late proteins from the RSD, which possess a kleptoplastid derived from *Phaeocystis antarctica*, resolve clearly with *Phaeocystis anatarctica* within the Phaeocystales, both considering single gene-trees (>70%) and the concatenated topology (Figures 1-2). We further noted a strong secondary signal to the family Isochrysidales in all kareniacean species including the RSD, even normalizing for dataset size (Figure S2), although many were specific to individual genera (Figure S3). Finally, we searched the new transcriptomes for contigs potentially representing plastid transcripts (recovering 13 to 39 genes in each case) and reconstructed a preliminary plastid phylogeny based concatenated nucleotide sequences. Consistent with previous studies, the fucoxanthin plastid genomes resolved as a long-branch sister group to all Prymnesiophyceae, although with a notable separation of *Takayama* from *Karenia* and *Karlodinium* (Figure S4) (Choi et al., 2017; Kawachi et al., 2021).

Finally, we used the whole transcriptomes as input for the PhyloFisher pipeline (Tice et al., 2021) to construct a matrix of 241 pan-eukaryotic nuclear-encoded genes to obtain a reference phylogeny of the studied kareniacean nuclear genomes in the context of full eukaryotic tree of life. The topology reconstructed from PhyloFisher matrix is consistent with recently published rRNA trees (Takahashi et al., 2019; Ok et al., 2021) with *Karenia* as sister to both *Karlodinium* and *Takayama* (Figure 2B), we note however that this phylogenetic topology differs to those reconstructed with both the plastid-late and plastid-encoded genes, which place *Karenia* and *Karlodinium* together, to the exclusion of *Takayama* (Figure 2; Figure S4). The monophyletic grouping of *Karlodinium* and *Takayama* was robustly rejected by approximately unbiased (AU) tests of the plastid-late and plastid-encoded tree topologies (p=0.006). The apparent incongruence between the placement of the *Takayama* plastid in the plastid-late, plastid-encoded and concatenated nuclear topologies raises questions concerning its endosymbiotic origin relative to other fucoxanthin-containing lineages.

### A hybrid plastid proteome with species-specific innovations

The plastid proteins of each kareniacean were automatically annotated with enzymatic functions based mainly on KEGG annotations, presented as a pivot table (Table S4). Considering χ^2^ enrichments of individual KEGG annotations (KO IDs) across all seven fucoxanthin-containing species, we noted a statistically significant enrichment (p < 0.01) in the plastid-late signal in the BRITE category “Photosynthesis”, combined category “Translation”, and KEGG Pathway “Porphyrin and chlorophyll metabolism”, while the plastid-early signal was enriched in BRITE categories “Membrane trafficking”, “Protein kinases, phosphatases, and associated proteins”, and “Ion channels”. Reflecting the probable plastid-early origin of the fucoxanthin plastid RNA processing machinery, categories relating to transcription and mRNA are not biased in their origin.

When analysed manually and qualitatively, additional evolutionary patterns were apparent in the overall metabolic map (Figure 3), some of which were previously described in *K. brevis* and *K. micrum* (Matsuo and Inagaki, 2018). These include notable differences between the genera and species in some pathways or enzymes, and several general particularities such as unexpectedly missing, duplicated, or laterally gained proteins, with full details supplied in Text S6 - Supplementary results and discussion.

**Figure 3:**
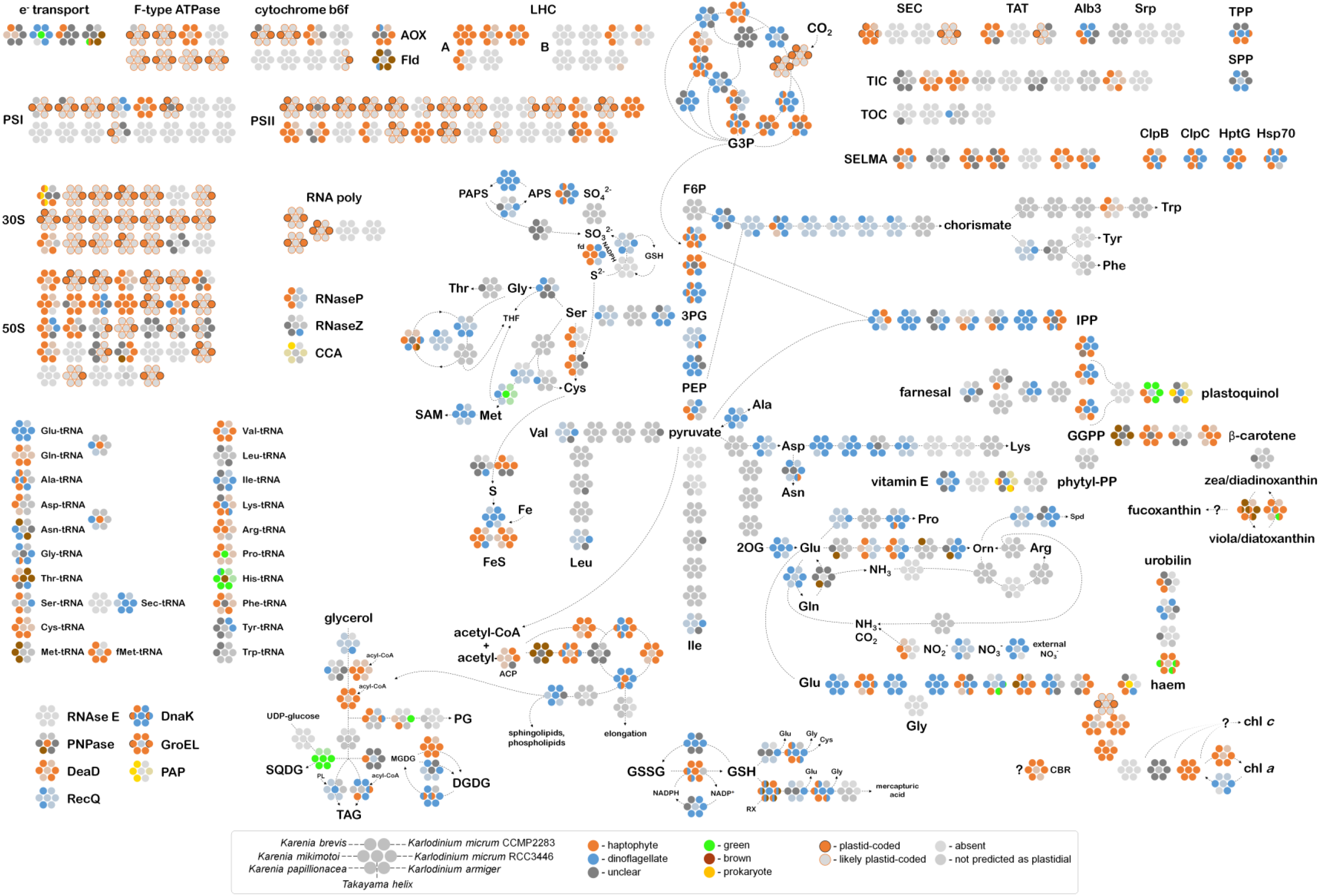
Reconstruction of major metabolic pathways of plastids of the seven kareniaceans, adapted from KEGG (Kanehisa and Goto, 2000). Each rosette (described in the legend) relates to the homologues of a specific nucleus-encoded and plastid-targeted protein, with each circle within each rosette corresponding to a different species, and different colours corresponding to different evolutionary affiliations. Proteins encoded in plastid genomes are provided for the incomplete genome assembly for *K. micrum* RCC3446, transcriptome-based set for *K. mikimotoi* and several sequences for *K. brevis*. A detailed breakdown of the individual sequences with their predicted origins used for this reconstruction are available in Table S3.

Reflecting KEGG enrichments, photosystems, light-harvesting complexes, photosynthetic electron transport chain and ATP synthase subunits are almost exclusively of plastid-late origin. So are ribosomal proteins and identified components of the protein import pathways, i.e., the “protein-level” housekeeping of the newest organelle, while the transcription machinery (when not directly plastid-coded, like RNA polymerase) is of mixed origin (see above). Furthermore, many of the pathways closely associated with photosynthetic function (i.e., syntheses of photosynthetic or photoprotective pigments) are of plastid-late origin, whereas the backbone pathways necessary for their synthesis (namely the tetrapyrrole pathway, preceding chlorophyll synthesis, and the MEP/DOXP pathway for synthesis of the IPP precursor of carotenoids) are of mixed or mostly plastid-early origin. Notably, we found almost no plastidial pathway to be of purely plastid-early origin, except perhaps the partial synthesis of some amino acids (Ala, Asp, Lys).

We report small numbers of conserved LGTs from other algal groups or bacteria, which are particularly concentrated among amino acyl-tRNA synthetases (HARS, TARS, MARS, NARS, PARS; see Figures S5.1-2 for selected exemplary trees), alongside with specific components of individual biosynthetic pathways (green SQD2, HST, haem oxygenase, and brown acetyl-CoA carboxylase, 15-cis-phytoene synthase, zeaxanthin epoxidase; exemplary trees in Figures S5.3-5). These genes were frequently detected across all fucoxanthin genera with strong bootstrap support and are therefore unlikely to result from contamination in individual transcriptomes or phylogenetic artifact.

Some otherwise broadly conserved proteins were not detected in any kareniacean transcriptomes, most notably SQD1, one of the two enzymes for the synthesis of essential plastidial glycerolipid sulfoquinovosyldiacylglycerol (SQDG), and the delta subunit of the photosynthetic ATP synthase (atpH, K02113). The former may be due to a very low expression level as it was previously missed in various algae even in dedicated studies (Riccio et al., 2020). The latter is more baffling as this conserved and essential subunit is usually plastid-encoded but was not retrieved as such in the previous plastid genome analyses (Gabrielsen et al., 2011; Dorrell et al., 2016) and was not identified in any of the full transcriptomes either. However, an unrelated and structurally dissimilar but functionally analogous subunit of mitochondrial ATPase (ATP5D, K02134) seems to be duplicated and retargeted to the plastid (Figure S6.4.1) and it is possible that this is in fact related to the absence of the gamma subunit.

### Targeting sequences of kareniacean plastid proteins vary with their evolutionary origin

Next, we wished to test if there are differences in the plastid targeting pre-sequences of plastid-late and plastid-early proteins, by comparison to analogous protein regions from peridinin dinoflagellate and haptophyte references. The signal peptides of kareniacean plastid-targeted proteins were found to typically contain a central LACLAC motif and a terminal GHG motif directly preceding the cleavage site regardless of whether they were of plastid-late or plastid-early origin (Figure 4A-B), whereas these motifs do not occur in haptophytes nor peridinin dinoflagellates (Figure 4C-D). Comparisons of the unique three-letter motifs associated with the plastid-targeting sequences of each group revealed particularly strong enrichments in the LACLAC and GHG motif and its variants in kareniacean proteins of plastid-late origin, compared to proteins of plastid-early origin and haptophyte and dinoflagellate equivalents (Figure 4E), despite similar overall amino acid composition of signal peptides across all datasets (Figure S7). In contrast, limited similarities were found in the transit peptides of kareniacean plastid-targeted proteins, apart from the double-arginine motif immediately following the cleavage site and conserved proline at +15 and arginine at +22 after the cleavage sites (Figure 4F).

**Figure 4:**
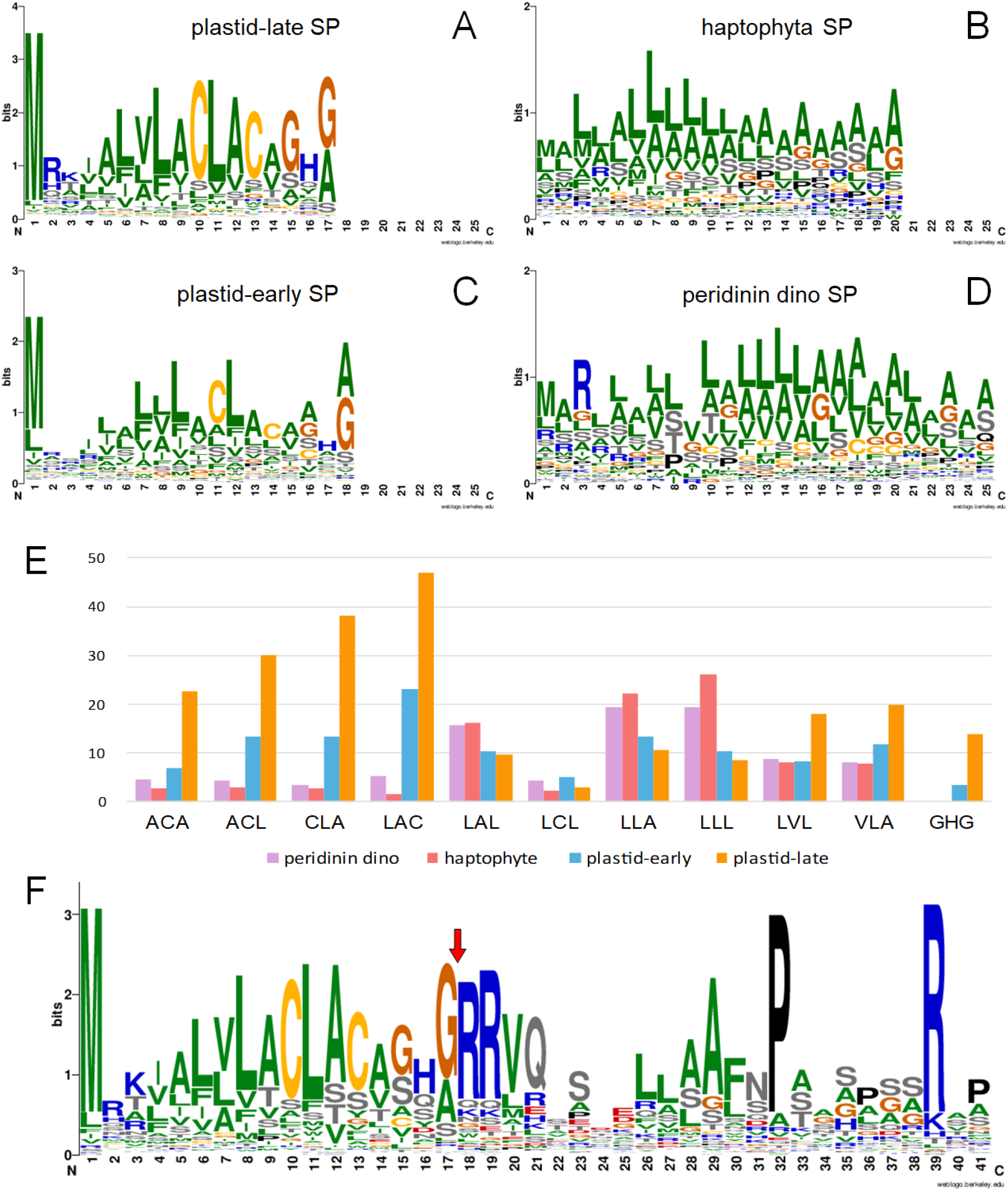
Sequence logos for aligned signal peptides of the four datasets of different evolutionary identity: haptophyte, peridinin dinoflagellate, plastid-late and plastid-early kareniacean (A-D), occurences of three-letter sequence motifs in signal peptides of each dataset, represented as a percentage of signal peptides in which at least one such motif occurs (E), and sequence logo of the signal peptide and partial transit peptide for kareniacean plastidial proteins regardless of their phylogenetic origin with red arrow denoting the signal peptide cleavage site (F).

### Global distribution of fucoxanthin dinoflagellates reveals *Karlodinium* as an outlier and a negative co-occurrence with their haptophyte progenitors

To provide biogeographical and ecological context for the interpretation of our transcriptomic and phylogenetic results we investigated the distribution of *Karenia*, *Karlodinium*, and *Takayama* in the *Tara* Oceans database using V9 18S metabarcoding (data available in Tables S5-8). Our analysis shows that *Karlodinium* clearly differs from the other two genera in its higher overall abundance and station occupancy and more cosmopolitan distribution, which is noticeable especially in the Indian and Arctic Oceans (Figure 5). Partial Least Square (PLS) analysis shows a set of environmental variables (salinity, silicate, iron) positively correlated with abundances of both *Karenia* and *Takayma* and also haptophytes as a whole, but at the same time negatively correlated to *Karlodinium* (Figure S8), further illustrating that the latter genus is quite distant from the rest in its biogeographical pattern. There was no correlation between any of the three studied genera themselves (Figure S9).

**Figure 5:**
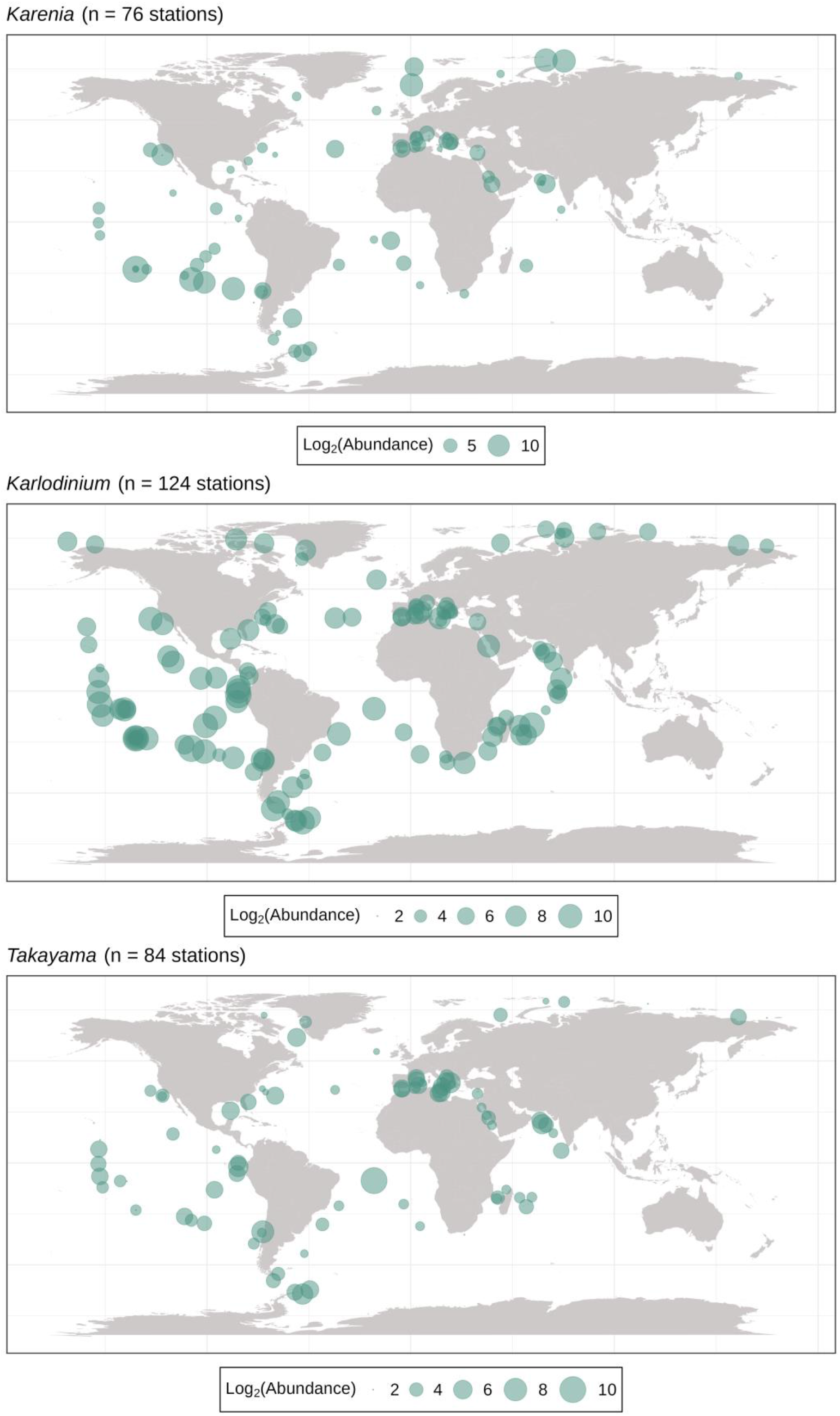
Abundances of the three studied kareniacean genera in Tara Oceans stations based on ribosomal 18S SSU V9 metabarcoding – *Karlodinium* exhibits higher global abundance and more cosmopolitan distribution in comparison to Karenia and *Takayama*; for a correlation circle with individual environmental variables, see Figure S9; none of the genera display significant positive or negative correlation to one another (Figure S10).

We further performed PLS analysis of the abundances of the three kareniacean genera and haptophytes, divided by family. We note an overall negative correlation to Chrysochromulinales, Phaeocystales, and Coccolithales, and positive correlation to Pavlovales (Figure 6), reflected by the overall preference of fucoxanthin dinoflagellates for temperate and tropical stations. We note that the negative association of present-day fucoxanthin dinoflagellates with Chrysochromulinales contrasts with the probable Chrysochromulinalean signal within their plastid proteomes.

**Figure 6:**
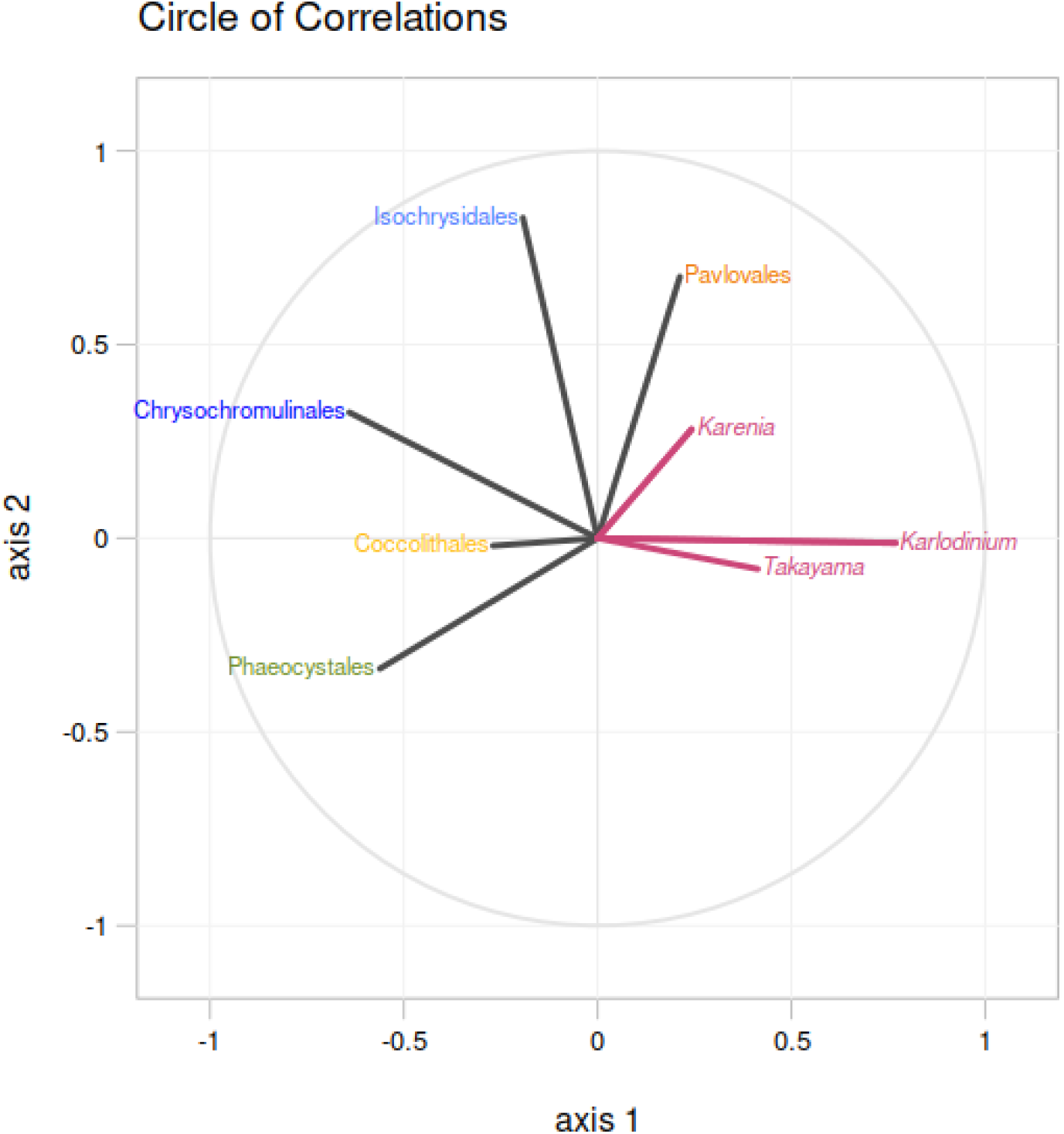
Correlation circle based on the Partial Least Square analysis of the *Tara* Oceans abundances (surface depth only) of the three studied kareniacean genera against those of different haptophyte families (colour-coding is the same as in Figure 1B) as observable variables; all three kareniaceans are negatively correlated to Chrysochromulinales, Coccolithales, and Phaeocystales but positively correlated to Pavlovales while only *Karenia* is positively correlated to Isochrysidales.

## Discussion

In this study, we harness novel and previously sequenced transcriptome data, customized *in silico* prediction, automated phylogenomics, and metagenomic analyses from *Tara* Oceans to reveal the proteomic composition and deeper taxonomic affinities of the fucoxanthin-containing plastids of the kareniacean dinoflagellates, a group of a special evolutionary importance as a tractable model system for studying the serial endosymbiotic acquisition of plastids. Thus, our results illuminate the mechanistics of a fundamental process that may underpin vast tracts of chloroplast evolution.

Using a combined phylogenetic matrix of 22 curated plastid-targeted proteins most closely related to haptophytes we were able to propose, to our knowledge for the first time, an origin for the fucoxanthin plastid origin with a specific haptophyte subgroup. Although the single-gene tree topologies vary greatly, often returning Kareniaceae as non-monophyletic, a plurality identify a probable origin within the Chrysochromulinales, and the refined combined phylogeny retrieves Kareniaceae as monophyletic and sister to Chrysochromulinales with maximum support. The non-negligible number of proteins exhibiting evolutionary affiliation with other groups / trees exhibiting conflicting topologies may be a result of either phylogenetic noise, or LGT from other haptophytes having facilitated the establishment of the fucoxanthin plastid in a taxonomically localized version of the “shopping bag model” (Larkum et al., 2007), particularly if the kareniacean ancestor was specialized to predation on haptophyte prey. We note that the divergence date inferred between Chrysochromulinales and Prymnesiales within the haptophytes from molecular clock data, i.e., between approximately 150 and 300 million years ago (Medlin et al., 2008; Liu et al., 2009), is coherent with the probable divergence of Kareniaceae from other dinoflagellate groups (no more than 250 million years ago), and is certainly more plausible than an origin between the divergence of the Pavlovophytes and radiation of Prymnesiophyceae (> 500 million years ago) (Parfrey et al., 2011; Strassert et al., 2021) as has been inferred from plastid-encoded gene phylogenies to demarcate the origin of the fucoxanthin plastid (Kawachi et al., 2021).

Our phylogenetic results are further correlated with environmental abundances in present-day metagenomic data. Chrysochromulinales, the presumed donor group of the plastid and also largest portion of the sortable haptophyte-like genes, is most strongly anti-correlated to all three kareniaceans, while the only group with which they are consistently positively correlated, Pavlovales, is the one that had the most negligible genetic imprint on their plastid proteins (compare Figure 1B and Figure 6). Isochrysidales, the genetic imprint of which seems to be the second largest, are positively correlated only to *Karenia*, not *Karlodinium* and *Takayama*. This pattern might reflect an active predation avoidance that may have evolved in those haptophyte lineages that were ancestrally common prey and therefore also potential gene and plastid source for kareniaceans, while the very weak correlation of Pavlovales to kareniacean dinoflagellates suggests no or very little predator-prey interaction, reflected in the limited contribution of the Pavlovales to the fucoxanthin plastid proteome. Nonetheless, this result is not easy to interpret without more data regarding the current predator-prey intractions between kareniaceans and haptophytes and their specificity, e.g., whether this may point towards predator avoidance and/or adaptation for different niches or, conversely, efficient clearing of the prey population by the predators Alternatively, the different geographical distributions of the chrysochromulinalean and kareniacean lineages may reflect contrasting environmental preferences. In this case, the acquisition of the fucoxanthin plastid by the kareniacean dinoflagellate host may have been accompanied by a transition to a new ecological niche, i.e., into tropical and oligotrophic instead of polar and nutrient-rich waters (Penot et al., 2022), with the associated physiological constraints directing its subsequent evolution. It is important to note in either case that the current biogeographical patterns of each lineage may not be at all indicative of the situation at time of the endosymbiotic event. The exact significance of the ecological interactions between the progenitors of Kareniaceae, and the haptophyte ancestors of their fucoxanthin plastid, may be best inferred via ancestral niche reconstruction for each lineage (as e.g., have been attempted by Moen and Morlon, 2014).

Strikingly, the plastid protein evolution suggested by our data - *Karlodinium* as sister to *Karenia* instead of *Takayama* – does not copy the nuclear topology of the kareniacean lineage. We even note that the plastid-encoded gene topology produced does not necessarily suggest monophyly of the Kareniacean plastid genome. While we cannot exclude that this is a result of gene duplication and differential retention of paralogs (Qiu et al., 2012), limited phylogenetic signal, and/or independent horizontal acquisitions of plastid-targeted proteins by *Takayama*, our concatenated topology need not imply a truly singular evolutionary origin of the organelle. *Takayama* differs from other kareniaceans in the shape of the feeding apparatus and preferred prey size - it feeds on various other dinoflagellates including large species (over 60μm) (Jeong et al., 2016), and may have consumed different prey than other fucoxanthin-containing lineages, which in turn may have differentially biased the potential sources of posterior LGT. At the same time, recent SSU and LSU phylogenies of Kareniaceae incorporate multiple species that do not possess a fucoxanthin plastid, that is RSD (Hehenberger et al., 2019), *Gertia* (Takahashi et al., 2019), and *Shimiella* (Ok et al., 2021) as closer relatives to *Karlodinium* and *Takayama* to the consistent exclusion of *Karenia*. RSD and *Gertia* retain the ancestral perdinin plastid, in a residual and slightly reduced but photosynthetically active and peridinin-containing forms, respectively. Phylogenetic analyses of the RSD by us and other authors shows next to no haptophyte-like genes shared with the other three genera, strongly suggesting that it indeed never possessed a stable fucoxanthin plastid (Hehenberger et al., 2019). We propose a scenario in which the fucoxanthin plastid of kareniaceans is indeed monophyletic, but was first acquired by the specific lineage leading to *Karlodinium* and *Takayama* after the divergence of the lineage(s) leading to the extant representatives with no or residual peridinin plastids. After this plastid was stabilized in the lineage leading to *Karlodinium*, it may have undergone an additional round of serial endosymbiosis spreading to the common ancestor of *Karenia* (Figure 7). We note that predator-prey relationships are not uncommon even between two toxin-producing dinoflagellates (Jeong et al., 2016). Conversely, the smaller size (de Salas et al., 2008) and cosmopolitan distribution (Figure 5) of *Karlodinium* might make it a potential prey for *Karenia*, although this awaits direct observation. Alternatively, the hypothetical additional serial endosymbiosis may have involved *Takayama* and a close relative of the chrysochromulinalean plastid donor of *Karenia* and *Karlodinium*.

**Figure 7:**
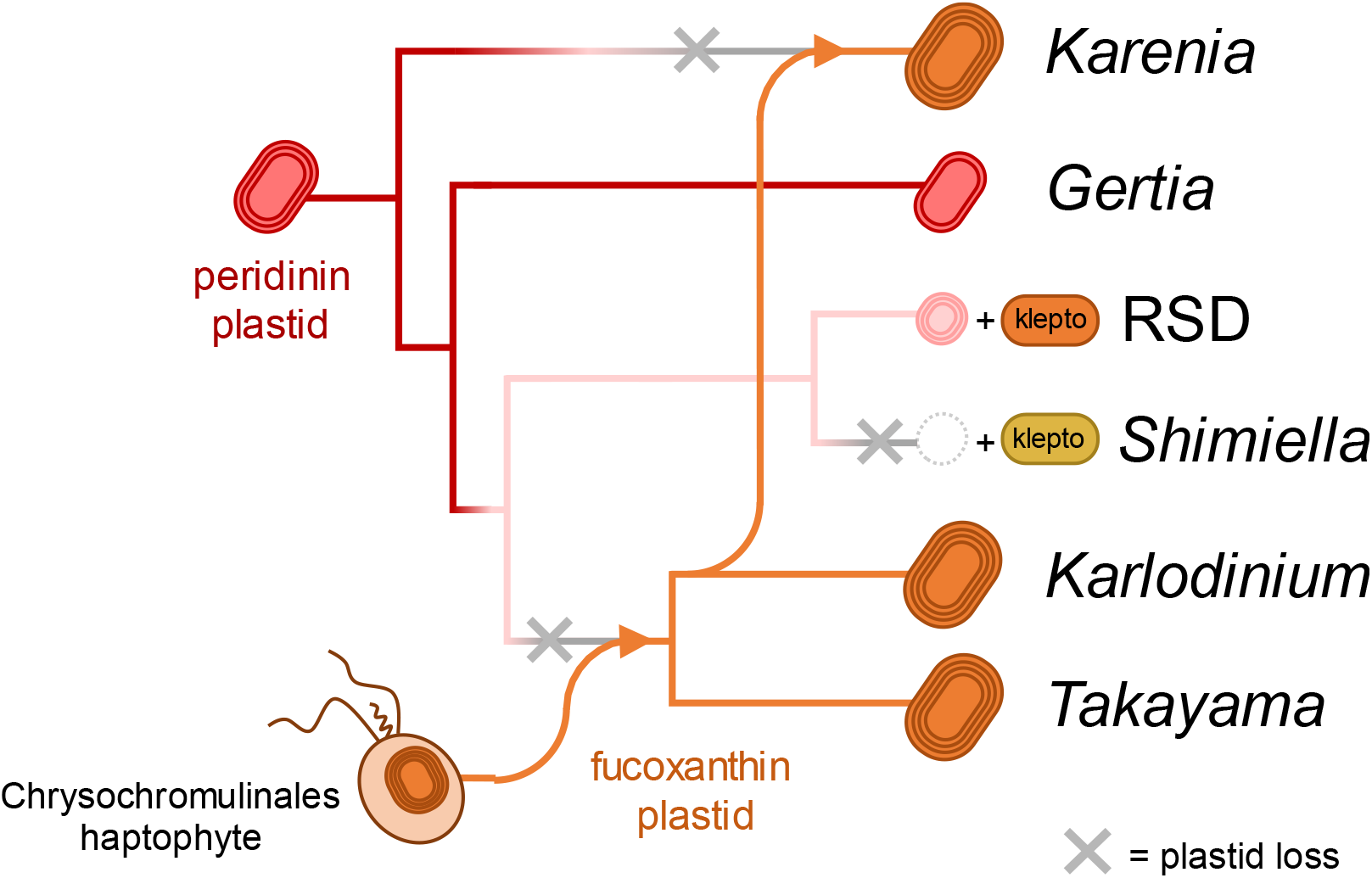
An evolutionary scenario involving an additional serial endosymbiosis between *Karlodinium*-like donor and an ancestor of *Karenia* which could explain the apparent non-monophyly of the fucoxanthin plastid suggested by the phylogenetic incongruence revealed by the placement of recently described lineages without the organelle (Takahashi et al., 2019; Ok et al., 2021) and reconstruction based on plastid-late proteins (see Figure 2). Plastid membrane layers and ultrastructures are drawn based on current microscopic observations for each species.

The evolutionary patterns in the compiled plastid metabolic pathways and complexes provides insights into the specific roles of EGT, LGT and host-derived proteins in the early evolution of the fucoxanthin plastid. The preferential incorporation of plastid-late pathways into photosynthesis-associated pathways may reflect that these functions were not encoded in the kareniacean host and that the new plastid (or much less frequently LGT) were the only available sources. In contrast, the retention of dinoflagellate-derived proteins in other pathways (e.g., isoprenoid, haem and amino acid biosynthesis) implies that the kareniacean ancestor retained these functions. We note that these functions are also often associated with protist lineages that have secondarily lost photosynthesis but retain vestigial plastids (Hadariová et al., 2018; Dorrell et al., 2019). It has previously been suggested for the RSD that the peridinin plastid is retained in a non-photosynthetic form (Hehenberger et al., 2019), and such an organelle might be the donor of the plastid-early pathways now found in the fucoxanthin plastid. One interesting case illustrating this scenario is the plastid-early chlorophyllase, the only enzyme of this origin in the chlorophyll branch of tetrapyrrole metabolism, which might have been retained by a heterotrophic kareniacean ancestor for the purpose of phytol recycling from chlorophyll contained in its algal prey. Ultimately, however, the presence (or eventual fate) of the peridinin-containing plastid in fucoxanthin-containing lineages awaits structural confirmation via microscopy.

The predominantly plastid-late origins of plastid translation but apparent retention of ancestral RNA processing pathways (e.g., poly(U) tail addition and substitutional editing) in the fucoxanthin plastid is puzzling in this regard, insofar as the peridinin plastid only encodes core photosystem subunits, and a non-photosynthetic plastid would be expected to lose its plastid genome (Dorrell and Howe, 2012; Jackson et al., 2013; Cooney et al., 2022). We note that RNA editing is widespread however in dinoflagellate mitochondrial and nuclear genomes (Lin et al., 2002; Liew et al., 2017), and also present in the green plastids of *Lepidodinium* that also secondarily replaced the peridinin-containing ones (Matsuo et al., 2022), and it remains to be determined if the fucoxanthin plastid RNA processing machinery is of relict peridinin plastid or non-plastidial host origin. It is also possible that there has been a direct niche competition between the peridinin and fucoxanthin plastid that may have coexisted in the same host for a period of time with possibly different selective pressure on retention of their respective proteins based on their interaction with plastid-encoded components, e.g., extrinsic photosystem subunits not assembling correctly with their intrinsic haptophyte-like counterparts. This would explain the strong bias towards plastid-late signal in photosynthetic machinery and ribosomes without the need for a period of heterotrophy with non-photosynthesizing plastid in the scenario, which could in turn make the abovementioned retention of peridinin-containing plastid in *Gertia* less baffling.

The general evolutionary division in the plastid metabolic map is not between plastid-late and plastid-early components but rather between parts strongly biased towards plastid-late and parts with no bias, assembled from both sources, likely in an evolutionarily neutral, patchwork-like way. This patchwork pattern does not only manifest at the level of one particular pathway being formed by enzymes of different evolutionary affiliations but often also at the level of one step in the pathway being carried out by homologs of different origin in different organisms. This division sometimes goes between the genera (e.g., plastid-late in *Karenia*, plastid-early in *Karlodinium*, consistent with the abovementioned overall higher proportion of plastid-late signal in the former; or *Takayama* appearing different from the rest) but sometimes even between individual species with no clear pattern. We propose that fucoxanthin plastid establishment may have been a relatively relaxed and gradual process during which peridinin-like and haptophyte-like genes may have both been present and competed for place in the nascent organelle (which was ultimately determined by neutral selection), not only during initial stages but also after the divergence of the genera and even species. We note that this phenomenon is also reflected in the partial plastid genomes assembled from *Karenia* and *Karlodinium*, which contain partially non-overlapping sets of genes that suggest independent post-endosymbiotic plastid genome reduction in each lineage (Gabrielsen et al., 2011; Richardson et al., 2014; Dorrell et al., 2016). It remains to be understood why a higher proportion of plastid-late proteins are observed in *Karenia* than *Karlodinium* or *Takayama*, which may variously relate to a greater retention of plastid-early genes in the lineage leading to *Karlodinium* and *Takayama*, greater post-endosymbiotic reduction and relocation of plastid-encoded genes from the *Karenia* plastid to the nucleus, greater paralogizations of plastid-late genes, or potentially additional LGT of genes from haptophytes specifically into the *Karenia* lineage. Deeper sampling of fucoxanthin plastid genomes, and of the haptophyte lineages they predate, may provide clues into the mechanistic processes guiding their deep evolution.

From the diversity of signal sequences targeting proteins into fucoxanthin plastids, we consider possible structural similarities that may reveal their underlying evolutionary origin. The most striking feature of kareniacean signal peptides (both of plastid-late and plastid-early proteins) is their high degree of conservation, which is evident from the comparisons of sequence logos (Figure 4A-D) and the fact that they are highly similar to one another, suggesting either their shared origin or very strict evolutionary pressure on their sequence. At the same time, they are very dissimilar to those of either of their potential sources, peridinin dinoflagellates and haptophytes. If they were indeed inherited from one or the other, they evolved profoundly under very strong selection to the point of being more strictly conserved than their ancestors and untraceable. The other possibility is that they arose *de novo* during fucoxanthin plastid establishment, and their highly similar sequence reflects their common and possibly relatively recent origin and spread throughout the kareniacean genome. Previous studies have suggested the significance of transposable elements for the recruitment of targeting sequences to nascent nucleus-encoded and plastid-targeted proteins (Burki et al., 2012), and the recent assembly of a draft *Karenia brevis* nuclear genome may allow us to assess to what extent transposons were involved in the transition of a transient kareniacean symbiont to an integrated organelle whose biogenesis is controlled by the host and its nuclear genome (Nelson et al., 2021).

Finally, we note that the basic metabarcoding analysis of the biogeographical distribution of Kareniaceae in *Tara* Oceans data is preliminary, particularly as it relies on analysis for the three genera due to the lack of species-level assignations of individual meta-barcodes. Meta-genome and meta-transcriptome abundances of plastid-late genes may enable a more precise enumeration of the environmental presence of the fucoxanthin plastid, particular by allowing the differentiation of fucoxanthin-containing Kareniaceae from peridinin-containing and non-photosynthetic relatives. Due to the almost completely translational regulation of the dinoflagellate nuclear genome (Lidie et al., 2005; Carradec et al., 2018), elucidating the precise physiological functions of individual fucoxanthin plastid proteins may however depend on more elaborate approaches, e.g., ribosome profiling of cultured species or environmental transcriptome data (Bowazolo et al., 2022). These eco-physiological insights may also be specific to each individual fucoxanthin dinoflagellate genus, particularly *Takayama* which seems to be notably separated from the rest in terms of its phylogenetic signal ratio and distribution among plastidial functions, as well as in its position in the phylogenies retrieved from plastid-targeted and plastid-coded genes; and *Karlodinium*, which represents an ecological outlier regarding abundance, stations occupancy, and correlation with environmental factors.

## Conclusions

Seven *in silico* plastid proteomes of fucoxanthin plastid-bearing kareniaceans were retrieved from already available and newly generated transcriptomic data and phylogenetically analysed using large-scale automated tree building and sorting. The ratio between plastid-late and plastid-early phylogenetic signal differs between the three studied genera and its uneven distribution among functionally annotated plastid enzymes in individual lineages suggests that the serial plastid establishment and formation of its evolutionarily mosaic proteome was a relatively long and relaxed process with room for significant diversification. Phylogenomic analyses of the haptophyte-like sequences reveal Chrysochromulinales as the closest relatives and possible donor lineage of the fucoxanthin plastid for the first time but at the same time suggests that its evolutionary history inside Kareniaceae was not straightforward and may have involved extensive horizontal gene flow or even additional serial endosymbiosis. Additionally, we reconstructed a comprehensive composite metabolic map of the fucoxanthin plastid, investigated kareniacean plastid signal sequences with regard to their phylogenetic origin, and analysed the global distribution of the three kareniacean genera in regard to both abiotic and biotic factors. We emphasise the importance of considering micro-evolutionary variation in the establishment of plastids in the Kareniacean dinoflagellates, and across the broader algal tree of life.

## Materials and methods

### Reference database preparation

An in-house protein database was used for homology searches throughout this project. This database consisted of 155 transcriptomic and genomic datasets fetched from NCBI (https://www.ncbi.nlm.nih.gov), 1KP (https://db.cngb.org/onekp/; Leebens-Mack et al., 2019) and decontaminated MMETSP (Keeling et al., 2014) as described and in Dorrell et al., 2021: 30 for eubacteria, 10 for archaea, and 115 for eukaryotes with sampling spanning most of the diversity with focus on plastid-bearing groups (100 datasets) and particularly enriched in dinoflagellate and haptophyte sequences (37 and 34 datasets, respectively). More details and list of used datasets is available in Table S2).

### Data acquisition and *de novo* transcriptome sequencing

The previously published peptide sequences for *Karenia brevis*, *Karlodinium micrum* (syn. *veneficum*) and Ross Sea Dinoflagellate (RSD) were downloaded from their respective public repositories (Keeling et al., 2014; Ryan et al., 2014, Hehenberger et al., 2019) and decontaminated as described above. In case of *K. brevis*, the four available assemblies from four clones were pooled together and treated as a single dataset with redundant sequences removed randomly.

*Karenia mikimotoi* RCC1513, *Karenia papilionacea* RCC6516 and *Karlodinium* sp. RCC3446 were obtained from Roscoff Culture Collection (https://roscoff-culture-collection.org), *Takayama helix* CCMP3082 was obtained from the Bigelow National Center for Marine Algae and Microbiota (https://ncma.bigelow.org), and *Karlodinium armiger* K0668 was obtained from the Norwegian Culture Collection of Algae (https://niva-cca.no). The cultures were grown in natural seawater (Horsey Mere seal colony, North Sea) supplemented with k/2 or enriched artificial seawater on a 12 hrs light (50μE): 12 hrs dark cycle at 19°C for 4 weeks, to late-exponential phase (1-2 million cells/ml). Cells were concentrated by centrifugation at 4000 rpm for 10 minutes, washed three times in sterile marine PBS (mPBS), and snap-frozen in liquid nitrogen. Total cellular RNA was extracted as previously described (Dorrell and Howe, 2012) using Trizol (Invivogen) and chloroform phase extraction, isopropanol precipitation, and resuspended in nuclease-free water (Qiagen). RNA integrity was confirmed by electrophoresis on ethidium bromide-stained agarose gels and quantified using a nanodrop photospectrometer. Two μg RNA was subsequently treated with 2U RNA-free DNAse (Promega) following the manufacturers’ instructions, precipitated with isopropanol, resuspended in nuclease-free water and quality-checked and quantified again.

For each species, 1 μg DNase-treated RNA was polyA-tail selected, and strand-specific libraries were prepared with TruSeq Stranded mRNA Library Prep kit (Illumina). The resulting libraries were sequenced using a Novaseq flow cell (Fasteris) with 100 bp paired reads, followed by in-house base-calling and basic Q30 enumeration (https://www.fasteris.com).

The obtained paired-end reads were further quality controlled using the RNA-QC-chain toolkit (Zhou et al., 2018) by filtering based on phred score with the threshold of 30; and GC content with the allowed range set to 42-60% based on the expected values for kareniacean dinoflagellates (Lidie et al., 2005), and by removing reads potentially representing rRNA contamination (default settings were used for this step). Filtered reads were then assembled by Trinity (Grabherr et al., 2011) with --NO_SEQTK and -- trimmomatic options and the resulting assembly was selected for the longest isoforms by the respective script from the Trinity toolkit. The quality and completeness of the transcriptomic datasets were assessed by TrinityStats and BUSCO (eukaryota_odb10, n:255) (Simão et al., 2015). The datasets were translated to proteins by TransDecoder (with default settings; https://github.com/TransDecoder/TransDecoder) and subsequently decontaminated by homology search using LAST (Kiełbasa et al., 2011) against the in-house database and removing sequences homologous to eubacterial, archaeal, or metazoan sequences with e-value of 1E-10 or lower. The translated datasets were used for further analyses alongside the previously published transcriptomes in their translated forms and are available at https://figshare.com/articles/dataset/full_protein_datasets_tar/21647771. The sequence data were deposited at NCBI under BioProject ID PRJNA788341.

### Plastid signal prediction

SignalP 5.0 (Almagro Armenteros et al., 2019) was used for prediction of N-terminal signal peptides as it was determined the version providing the most specific plastid prediction during preliminary optimization. Both original sequences and sequences cut to the first methionine were analysed to accommodate for potential translated UTRs or spliced leaders, and the results were pooled afterwards prioritizing the positive signal peptide prediction in case of different results. The obtained SignalP results were converted into a tabular format compatible with SignalP versions 3.0-4.1 for downstream applications with a custom script (SP5toSP4.py). For the second step predicting the rest of the bipartite targeting signal, a modified version of ASAFind (Gruber et al., 2015; Füssy et al., 2019) was used with a custom scoring matrix (modASAFind.py, Table S9) that reflects the differences in sequence surrounding the signal peptide cleavage site of kareniaceans (for instance, instead of the phenylalanine motif, multiple arginine residues are typically present). The matrix was prepared based on sequence logos prepared by Seq2Logo (Thomsen and Nielsen, 2012; Figure S10) from model datasets of *Karenia brevis* and *Karlodinium micrum* proteins determined as having exclusively plastidial, mitochondrial, nuclear, or endomembrane functions, e.g., photosynthetic core machinery proteins for plastidial functions, and identified from reciprocal BLAST against well-curated proteomic datasets from other organisms (Butterfield et al., 2013; Klinger et al., 2013; Dorrell et al., 2017; Bannerman et al., 2018; Beauchemin and Morse, 2018), the queries and retrieved model sequences are available in Tables S10-11. The modified script and custom scoring matrix were subsequently tested and optimized for sensitivity and specificity, and in the final version represented an improvement of 56.4 and 0.5 percentage points, respectively over unmodified ASAFind. The detection thresholds used with SignalP were also tested, with no apparent improvement in specificity/sensitivity tradeoffs over the default settings; as well as prediction by HECTAR and combination of PrediSI and ChloroP (Gschloessl et al., 2008; Dorrell et al., 2019; Ebenezer et al., 2019; Figure S11-12, Tables S12-16).

Additionally, we employed an alternative prediction, previously tested on the unrelated eukaryote *Euglena gracilis* combining PrediSI (Hiller et al., 2004) and ChloroP (Emanuelsson et al., 1999) and proteins judged negative by modified ASAFind but positive by this approach were included in the predicted plastid proteomes as well albeit with prefix “SPTP-” denoting the presence of an alternative signal.

The presence of large numbers of highly similar sequences caused by very recent paralog duplication and/or alternative splicing is common in dinoflagellates. To reduce this complexity for phylogenetic analyses, all datasets were additionally clustered by cd-hit with an identity threshold of 0.9. This was performed after signal prediction to avoid accidentally discarding an isoform/paralog with predictable targeting signal in favor of one without it.

### Phylogenetic analysis

Each of the *in silico* plastid proteome datasets was processed independently in order not to omit any variant signals in individual proteins. First, homologs were mined by LAST (Kiełbasa et al., 2011) search with threshold e-value 1E-10 against the in-house database plus all the fucoxanthin dinoflagellate transcriptomes. The best hit from all organisms in the in-house dataset were retained alongside the best five hits from each other fucoxanthin dinoflagellate library to account for potential paralogs in fucoxanthin dinoflagellate transcriptomes. Proteins with no hits whatsoever or with no homologs in any taxa other than dinoflagellates (Kareniaceae or other) were annotated as “lineage-specific” or “dinoflagellate-specific” respectively and their phylogeny was not investigated further. The remaining homolog sets were aligned by MAFFT (version 7.310, Mar 17 2017; Katoh and Standley, 2013) with default settings and trimmed by TrimAl (version 1.4, Dec 17 2013; Capella-Gutierrez et al., 2009) with – gappyout parameter. The trimmed alignments were then filtered by a custom python script that discarded sequences comprising of more than 75% gaps and then rejected alignments shorter than 100 positions or containing less than 10 taxa. The satisfactory alignments were then used for maximum-likelihood tree construction by IQ-TREE (multicore version 2.0.5, May 15 2020; Minh et al., 2020) with automatic model selection and 1000 ultrafast bootstraps. For graphical summary of this pipeline, see Figure S2. All trees are available at https://figshare.com/articles/dataset/all-automatically-built-trees_tar/21647768; in-house scripts used in the pipeline are available among supplementary files (scripts.tar); as well as the final predicted plastid protein datasets with their phylogenetic annotations (phylo-annotated_proteins_sets.tar).

TreeSorter, a freeware program written in Python (https://github.com/vanclcode/treesorter/) builds abstract tree structure from a tree file in nexus format, takes arguments defining sets of taxa and criteria based on those sets that determine what constitutes a valid subtree. Each tree is traversed and the highest bootstrap value of an edge (bipartition) that produces a subtree containing the taxon of interest and matching the defined criteria is reported. Criteria allow for further quantification of taxon sets, such that minimum and/or maximum number or proportion of taxa from a particular set per subtree. TreeSorter was used for large-scale analysis of the tree topologies and sorting and annotating the seed plastid proteins by their probable evolutionary origin into custom categories: haptophyte (plastid-late), dinoflagellate (plastid-early), alveolate (non-plastid), green (i.e., green algal, plant, or chlorarachniophyte; potential LGT or EGT), brown (i.e., ochrophyte; potential LGT or EGT), prokaryotic (likely LGT) as sequences of these evolutionary origins were noticed in these organisms in previous studies (Nosenko et al., 2006; Waller et al., 2006), and other/unresolved. In order to be assigned to one of the categories, the seed protein had to be either sister to or nested within a clade comprising exclusively members of said category and of at least 2 taxa. The bootstrap value of the highest-supported branch dividing such clade from the rest of the tree was then saved as the numerical score of this sorting result. In case the protein can be assigned to more than one category, the one with higher bootstrap was selected, if they have equal score, the protein is considered unresolved, with the exception of nested categories (such as dinoflagellates and alveolates) where the wider category is selected. Bootstrap scores of 75 were generally considered as the lower threshold in interpreting the evolutionary origins, however under specific circumstances (e.g. when the same protein was assigned the same origin with score ≥75 in other studied species) lower scoring results are reported as well.

The *in silico* plastid-targeted proteomes were automatically annotated by KAAS and major metabolic pathways were reconstructed, either using KEGG Mapper (https://www.genome.jp/kegg/mapper/reconstruct.html), or manually by targeted HMMER search for missing enzymes or enzymes without representation in KEGG database and/ or based on their conserved domains as detected by PfamScan (Finn et al., 2014).

### Phylogenomic analysis

A concatenated phylogenetic matrix for plastid proteins was prepared using 22 genes that passed our criteria of a) tree-based evidence of plastid-late origins, b) occurrence in at least 5 of the 7 organisms, and c) manually verified functional annotations and plastid-associated functions: 1-acyl-sn-glycerol-3-phosphate acyltransferase, 3-hydroxyacyl-ACP dehydratase, 3-oxoacyl-ACP synthase, ACP, cysteinyl-tRNA synthetase, digalactosyldiacylglycerol synthase, glutaminyl-tRNA synthetase, haem oxygenase, chlorophyll a synthase, chlorophyllide b reductase, lycopene beta cyclase, magnesium chelatase subunit H, magnesium-protoporphyrin O-methyltransferase, protochlorophyllide reductase, PsbO, PsbP, PsbU, RP-L1, RP-L13, SecA, SufC, and SufD. Preliminary single-gene trees were manually checked to remove paralogs, very long branches, and likely cases of LGT, if present; these trees with the removed taxa highlighted, as well as the original and trimmed datasets, and final matrix are available as supplementary files (plastid_protein_concatenation_files.tar). The phylogenetic matrix for non-plastid proteins was prepared using the standard workflow of the PhyloFisher toolkit (Tice et al., 2021), with the manual step of paralog annotation by parasorter, and additional removal of sequences containing >66% of gaps; the matrix constructor statistics, input metadata, and the matrix itself available as supplementary files (phylofisher_files.tar). Both concatenated trees were then constructed by IQ-TREE with LG+C60+F model for the plastid protein matrix and posterior mean site frequency (PMSF) model (LG+C60+F+G with a guide tree constructed with C20) for the non-plastid matrix (Wang et al., 2018). Additionally, a constrained tree search and subsequent approximately unbiased (AU) topology test were performed by IQ-TREE on the plastid protein matrix to verify the recovered inner topology of Kareniaceae.

### Signal peptide analysis

Four experimental datasets for this analysis consisted of two sets of plastid-targeted plastid-early (264) and plastid-late (437) proteins with assigned KEGG annotation with uniquely plastidial function and two analogous sets selected from MMETSP transcriptomes for five peridinin dinoflagellates (358) and five haptophytes (272) (details in Table S17) based on presence of a) detectable signal peptide and b) the same KEGG annotations to produce comparable datasets without functional bias. Signal peptide regions of these proteins were determined by SignalP 5.0 (the region of approximately 25 positions immediately following the predicted cleavage site was considered as putative partial transit peptide region). The regions were aligned by ClustalW, manually curated to keep their conserved part and used to create sequence logos for each dataset using WebLogo (https://weblogo.berkeley.edu/logo.cgi; Crooks et al., 2004). The overall amino-acid composition and number of occurrences of three-letter sequence motifs suggested by the logos in each dataset was also determined.

### Meta-barcoding analysis of Kareniaceae distributions

To explore the distribution of the three Kareniaceae genera, the V9 18S rDNA metabarcoding data from *Tara* Oceans corresponding to the genera *Karenia*, *Karlodinium* and *Takayama* were extracted from Ibarbalz et al., 2019 (available at https://zenodo.org/record/3768510#.Ye-xqpHMJzp). Overall, a total of 69 barcodes were extracted for *Karenia*, 76 for *Karlodinium* and 131 for *Takayama*. Their respective distributions (at least equal to three reads) in the eukaryotic-enriched size fractions (Pierella Karlusich et al., 2019) (0.8-5 μm; 5-20 μm; 20-180 μm and 180-2000 μm) and two depths (surface and deep-chlorophyll maximum) were pooled together and plotted using the R package ‘mapdata’ v2.3.0 (http://cran.nexr.com/web/packages/mapdata/index.html). The occupancy was estimated as the percentage of stations in which their respective barcodes were identified. A correlation analysis was conducted on the pooled Kareniaceae size fractions and depths, including a total of 75 stations in which at least one of the organisms was retrieved, with the R package ‘corrplot’ (https://github.com/taiyun/corrplot; Wei and Simko, 2021). Two final Partial Least Square analyses were implemented with the R package ‘plsdepot’ (https://github.com/gastonstat/plsdepot) to explore the correlation between: 1) kareniacean abundances and the abundance of the 5 haptophyte taxa Chrysochromulinaceae, Coccolithales, Isochrysidales, Pavlovophyceae and Phaeocystales (V9 18S rDNA metabarcoding data extracted as above); 2) global kareniacean and haptophyte (all haptophyte taxa pooled) abundances compared with a diverse set of environmental variables (median values of nutrient concentrations of iron, nitrate, silicate, phosphate; and oxygen, salinity, temperature, all extracted from the PANGAEA database; Ardyna et al., 2017; Guidi et al., 2017b, 2017a). Both the abundance and environmental data considered for the Partial Least Square analyses were computed only for the surface with all four size fraction abundances pooled.

## Supporting information

modASAFind

phylo-annotated_proteins_sets

phylofisher_files

plastid_coded_nt_concatenation_files

plastid_protein_concatenation_files

scripts

SP5toSP4

## Acknowledgements

RGD and ANV acknowledge supporting from a CNRS Momentum Fellowship awarded to RGD, 2019-2021, and an ANR JCJC (“PanArctica”, ANR-21-CE02-0014) awarded to RGD, 2021-2022. RGD further acknowledges an ANR t-ERC (“ChloroMosaic”), awarded 2022. CB acknowledges funding from the European Research Council (ERC) under the European Union’s Horizon 2020 research and innovation programme (Diatomic; grant agreement No. 835067), the French Government ‘Investissements d’Avenir’ programmes MEMO LIFE (ANR-10-LABX-54) and PSL* Research University (ANR-1253 11-IDEX-0001-02), and from the Agence Nationale de la Recherche (BrownCut; grant agreement ANR-19-CE20-0020). The authors thank Elisabeth Hehenberger for sharing translated and decontaminated RSD data and consultation regarding their analysis, Lukáš Novák for consultation regarding phylogenomics, and Emmanuel Alastra for aid in media substrate preparation. This article is contribution [will be provided at proof stage] to *Tara* Oceans.

## Notes

### Competing Interest Statement

The authors have declared no competing interest.

