## Supplementary figures and text for "Divergent and diversified proteome content across a serially acquired plastid lineage"

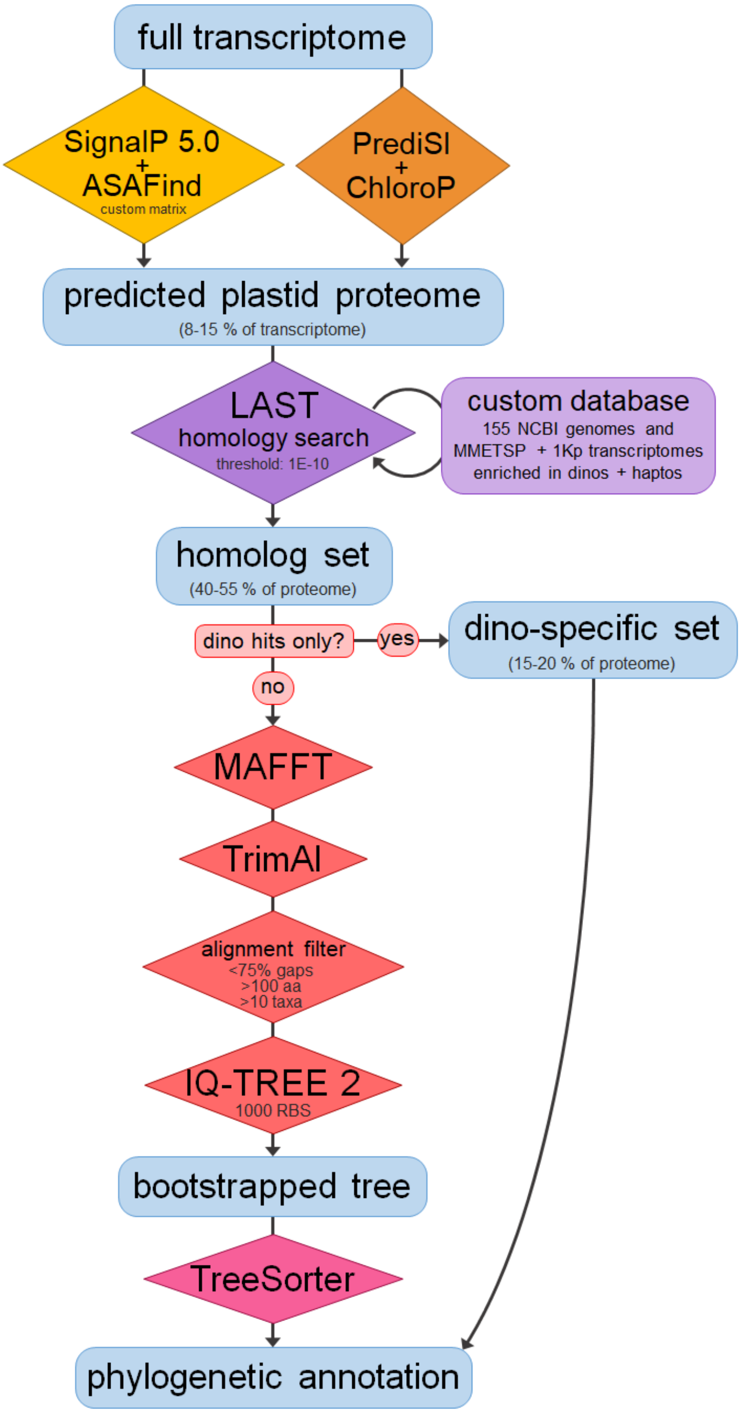


*Figure S1*: Graphical summary of the bioinformatic pipeline used to predict plastid-targeted proteins and obtain their phylogenetic annotations.

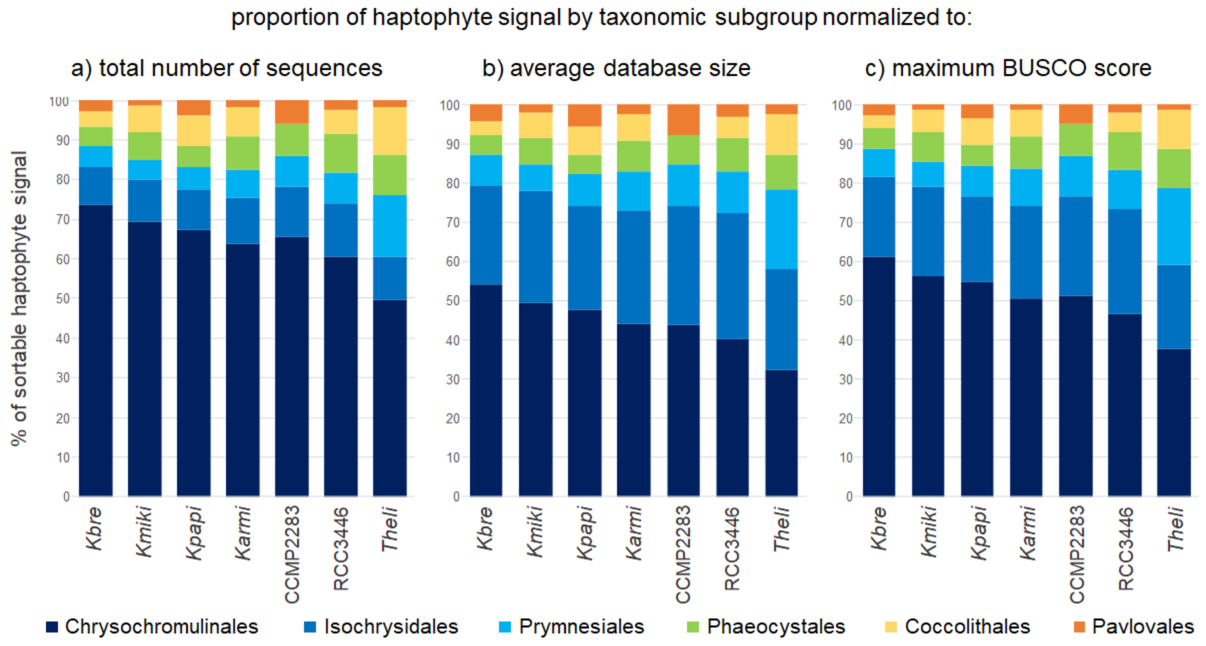


*Figure S2*: Ratios of the plastid-late proteins further sortable to respective haptophyte families normalized against sum of numbers of sequences in all the respective transcriptomes in our database (a), average size of these transcriptomes (b), and the maximum BUSCO score for each family (c).

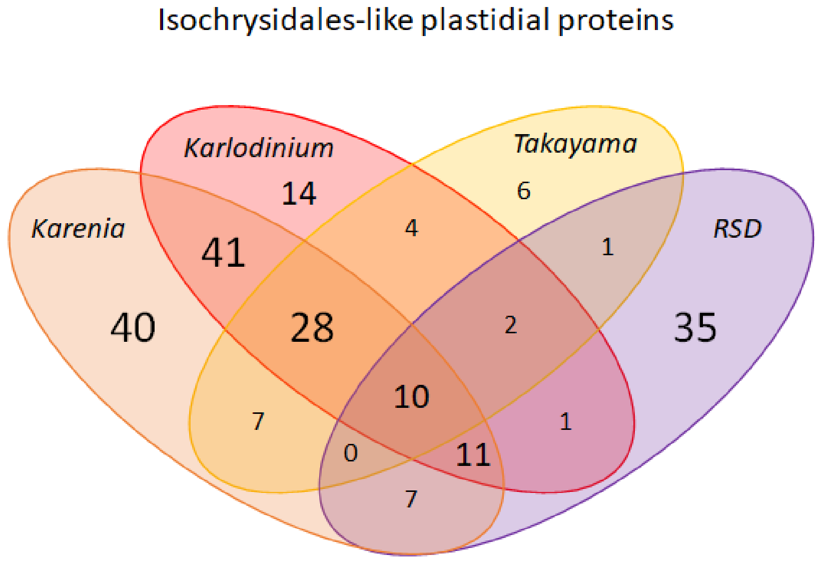


*Figure S3.1*: Venn diagram of the distribution of homologs of proteins with specific Isochrysidales-like origins recovered in fucoxanthin plastid proteomes; the number of shared homologs is the highest between *Karenia* and *Karlodinium* (41), followed by those shared by the three non-RSD genera (28), and only then those shared by all four (10). At the same time, the highest number of genus-specific Isochrysidales-like proteins are in *Karenia* (40) and RSD (35), and only in RSD, this number is higher than the total number of proteins shared with at least one other group. This relative isolation of RSD from the rest even in terms of LGT from other haptophyte groups reflect the different evolutionary origin of its plastid.

*Figure S3.2*: Single-gene tree generated by our pipeline with Karenia-brevis-SP1-CAMPEP-0189089492 (functionally annotated as acetylglutamate kinase) as a seed sequence showing the Isochrysidales-like phylogenetic affiliation of the retrieved *Karenia* homologs and separate, low-supported dinoflagellate-like origin of the *Karlodinium* homologs; a *Takayama* homolog was not identified for this protein while retrieved RSD homologs are *Phaeocystis*-like and bacterial-like. Bootstrap support is expressed by the branch colour (black for ≥90%, dark grey for ≥75%, light grey for <75%).

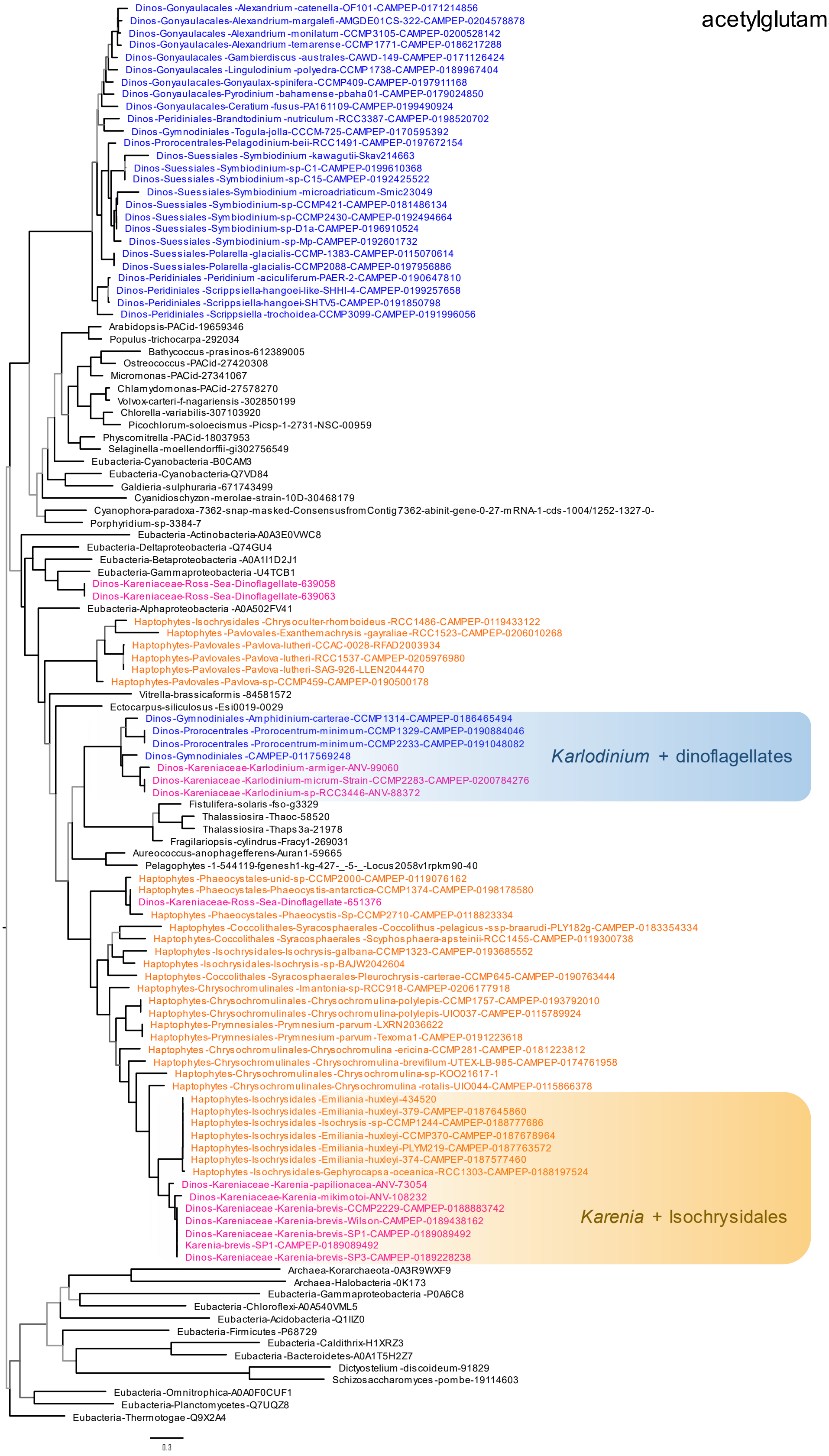

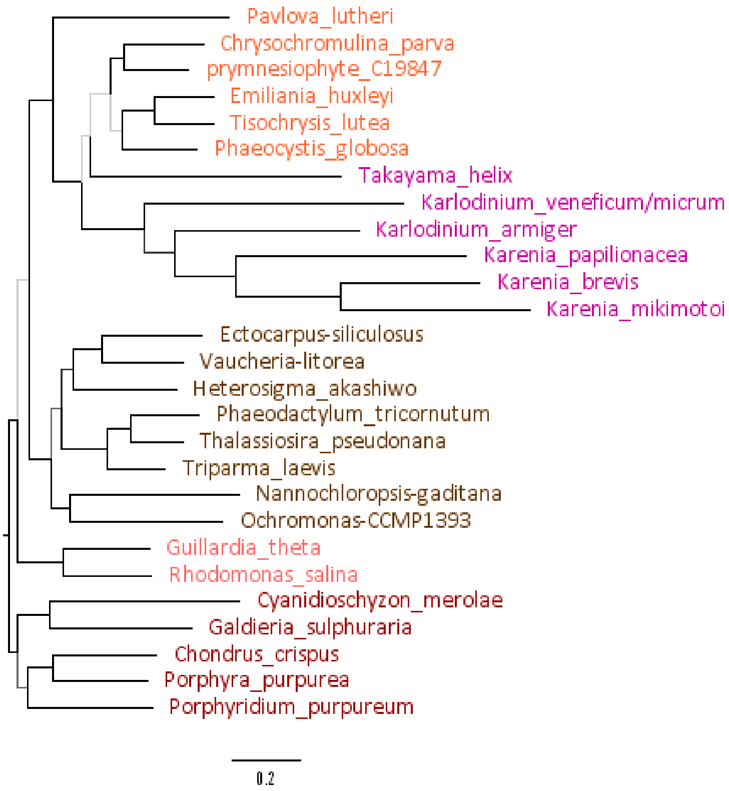


*Figure S4*: Tree reconstructed from a nucleotide phylogenetic matrix concatenated from 31 putatively plastidial trancripts (original datasets, alignments, and single-gene trees available in plastid_coded_nt_concatenation_files.tar); bootstrap support is expressed by the branch colour (black for ≥90%, dark grey for ≥75%, light grey for <75%). Kareniaceae appear as a long-branching paraphylum to the Prymnesiophyceae in this topology with *Takayama* split from the rest on an unsupported branch, and *Karenia* and *Karlodinium* forming a well-supported monophylum, and *Karlodinium* itself resolving as paraphyletic.

*Figures S5.1-5*: Selected phylogenetic trees of plastidial proteins putatively originated by LGT from green and brown algae; all trees are unrooted and constructed by the same pipeline as the rest of single-gene trees using all retrieved plastid-targeted kareniacean homologs as seed sequences before removing redundant hits. Bootstrap support is expressed by the branch colour (black for ≥90%, dark grey for ≥75%, light grey for <75%); sequences from the studied kareniaceans are coloured pink with proteins with predicted plastid-targeting signal in darker shade and annotated “CP” at the end of the ID; colour-coding for other dinoflagellates and haptophytes remains blue and orange, respectively, while green and brown algae (i.e. Chloroplastida and Ochrophyta) are highlighted in their respective colours.

*Figure S5.1*: Green origin of plastidial histidyl-tRNA synthetases of *Karenia* and *Karlodinium* and brown origin of the plastidial homolog in *Takayama*. A green-like homolog present in *Takayama* is not predicted as plastidial. The second kareniacean clade inside other dinoflagellates likely represents the mitochondrial or cytosolic form of the enzyme.

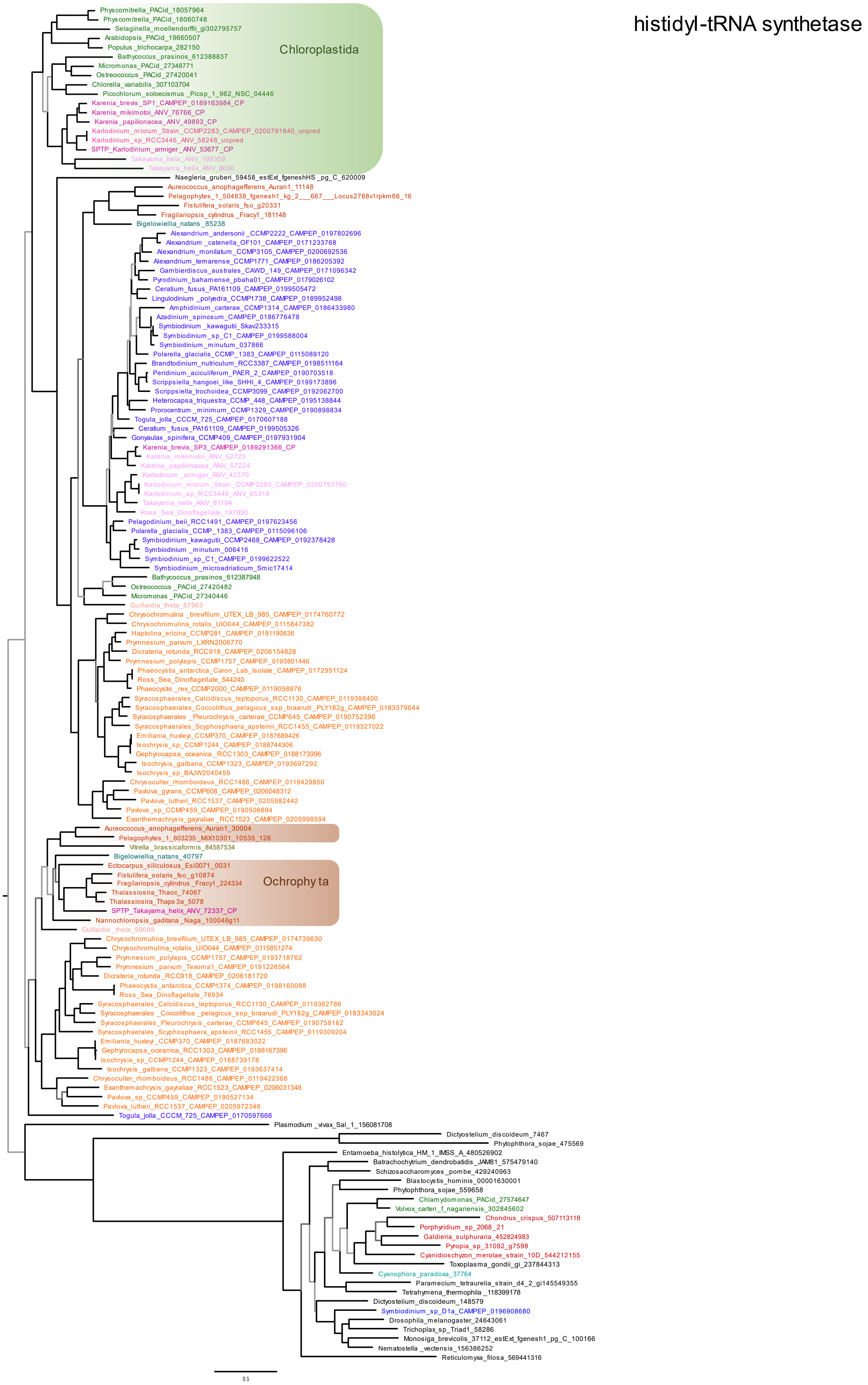

*Figure S5.2*: Brown origin of plastidial threonyl-tRNA synthetase present in representatives of all three genera.

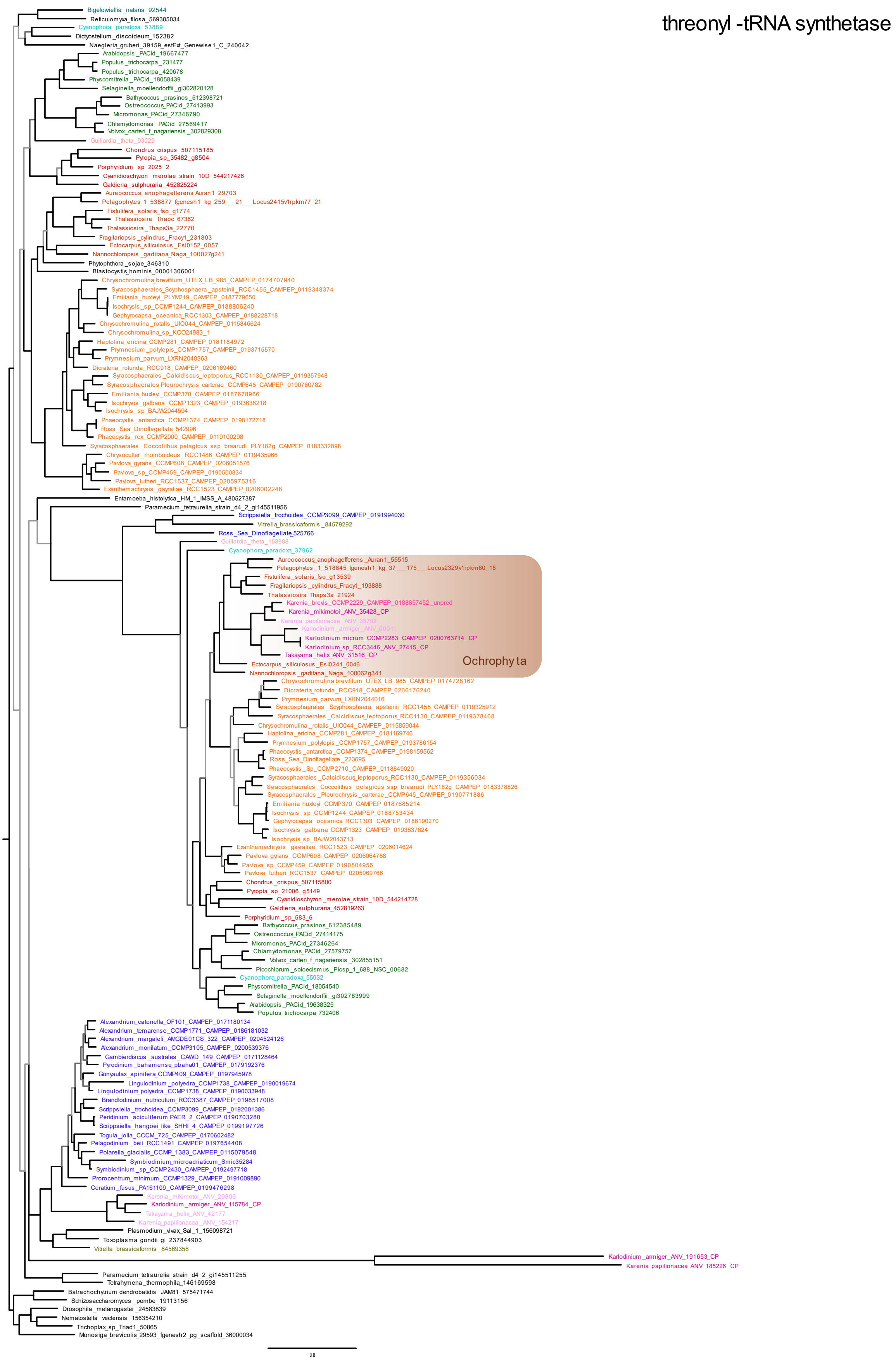

*Figure S5.3*: Green origin of sulfoquinovosyltransferase (SQD2) in all three genera.

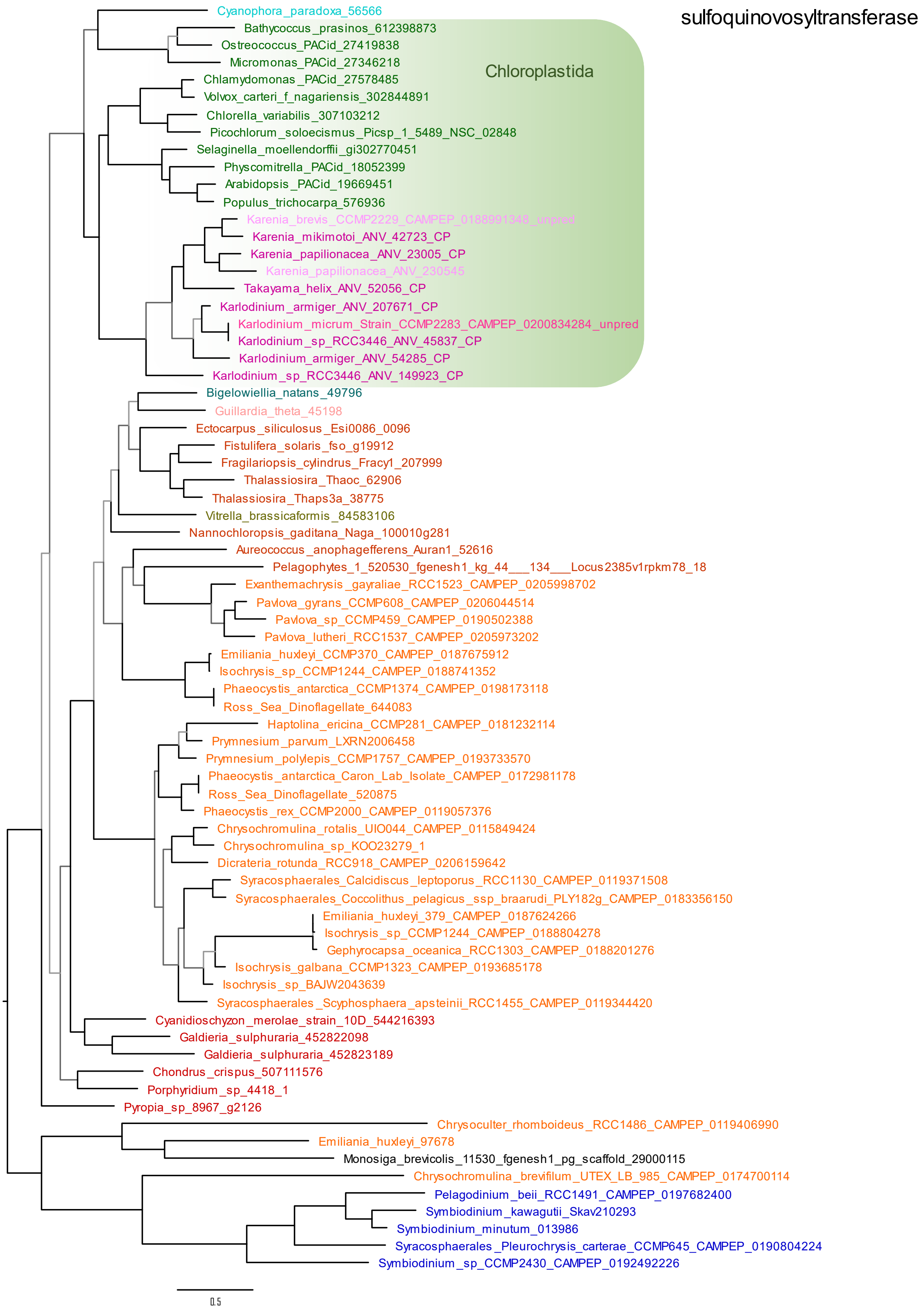

*Figure S5.4*: Green origin of one form of haem oxygenase in *Karenia*, *Karlodinium*, and the RSD (*Takayama* homolog was not retrieved); the second form is of haptophyte-like plastid-late origin in all genera.

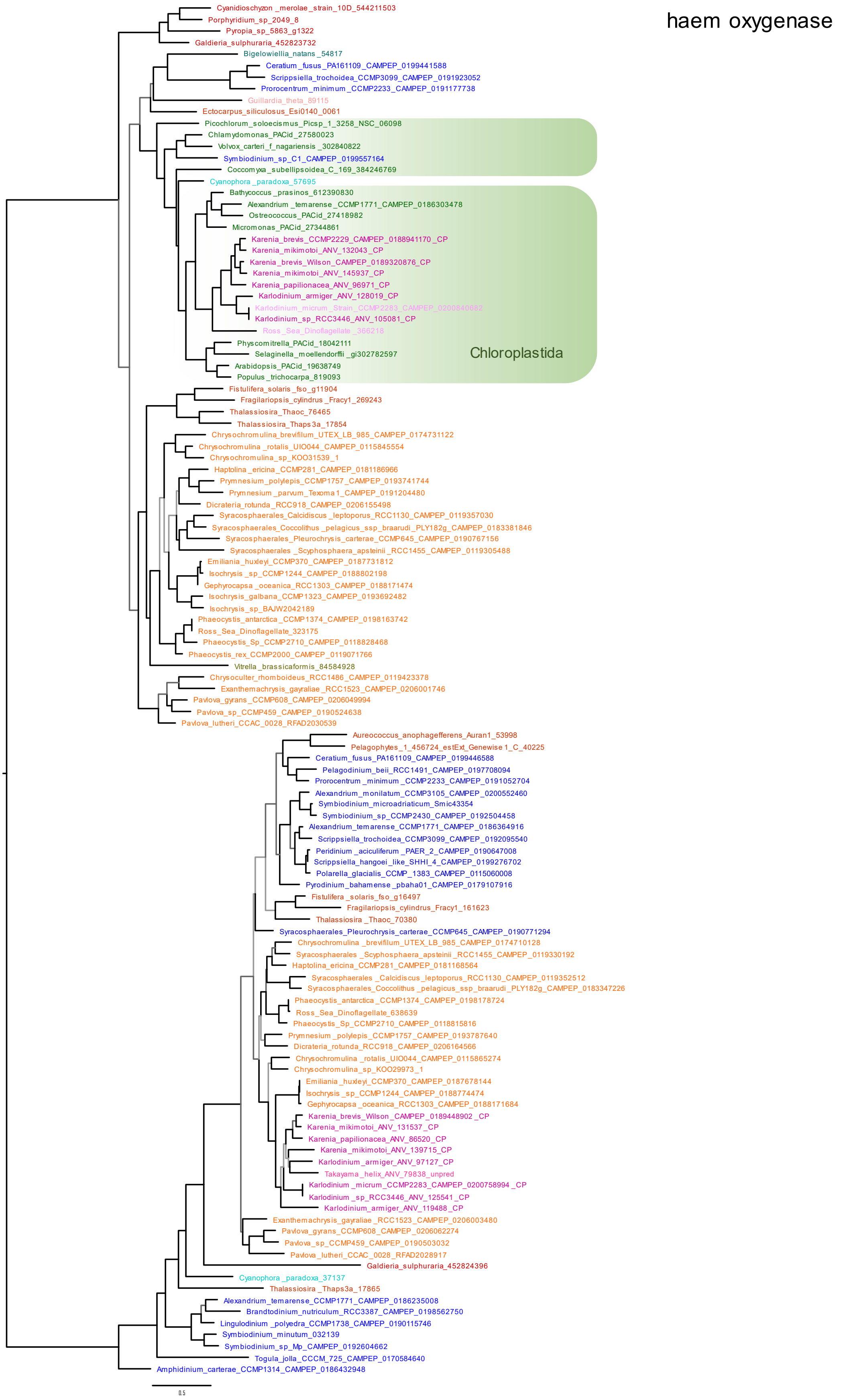

*Figure S5.5*: Brown origin of the 15-cis-phytoene synthase in all genera.

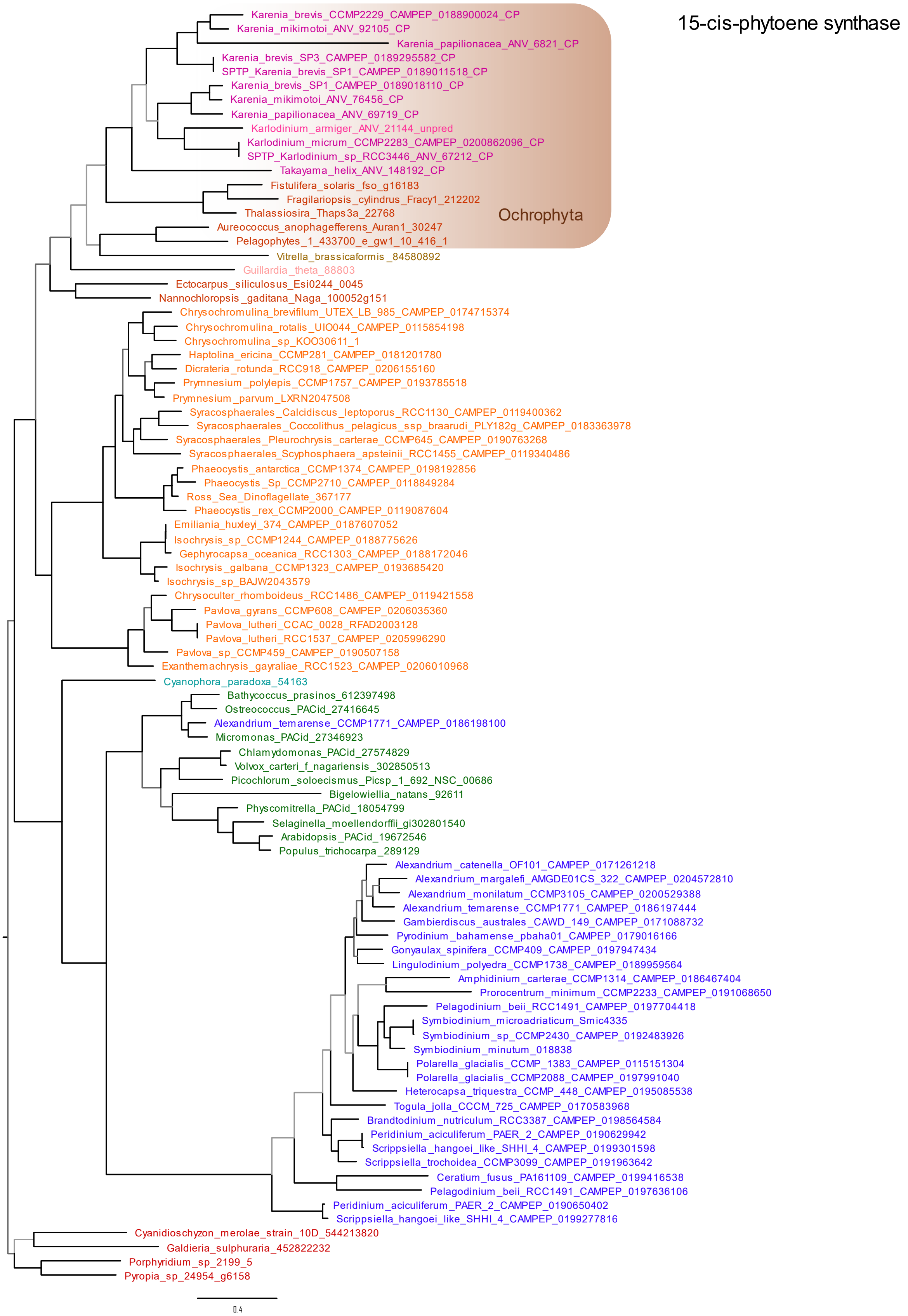

### *S6: Supplementary results and discussion: Plastidial metabolic pathways and their notable features*

#### *S6.1: Tetrapyrrole metabolism*

The tetrapyrrole pathway is very clearly divided into three parts: a backbone leading from glutamine to protoporphyrin, the haem and bilin pathway, and the chlorophyll-synthesizing branch (Tanaka and Tanaka, 2006). While the former two are slightly skewed towards a plastid-early signal but are generally of mixed origin (including isolated cases of putative LGT), the latter comprises almost exclusively of plastid-late proteins The only plastid-early enzyme associated specifically with chlorophyll is chlorophyllase, which has additional roles in phytol recycling to chlorophyll synthesis.

Two evolutionarily distinct versions of the biliverdin-producing haem oxygenase seem to be present in the plastid of representatives of both *Karenia* and *Karlodonium*: one of plastid-late and one of green-like origin (each in one or two copies, Figure S5.4). This enzyme is often present in multiple copies in photosynthetic organisms and it has been theorized that the copies may play slightly different roles in other reactions of bilin metabolism, especially in photosynthetic organisms where these molecules serve as chromophores of phytochromes  (Dammeyer and Frankenberg-Dinkel, 2008). The specific functions metabolic of the green-like and haptophyte-like haem oxygenases in the fucoxanthin plastid await experimental characterisation.

#### *S6.2: Terpenoid metabolism*

A similar break in the evolutionary pattern of the pathway can be observed between the backbone of terpenoid synthesis (MEP-DOXP pathway), which is mixed and often not consistent between the genera or even species, but overall slightly skewed towards plastid-early genes; and other pathways branching from it, most notably carotenoid metabolism (Coesel et al., 2008) which comprises mostly of plastid-late proteins and does not contain any plastid-early proteins. Two proteins (15-cis-phytoene synthase (Figure S5.5) and zeaxanthin epoxidase) were potentially gained laterally from brown algae.

The shorter branching pathways leading to plastoquinol and tocopherol contain two proteins likely gained laterally from bacteria (MPBQ/MSBQ methyltransferase / VTE3) and green algae (homogentisate solanesyltransferase / HST) but neither of the pathways seems to be complete (all-trans-nonaprenyl-diphosphate synthase and VTE1 were not identified by HMMER search in any of the transcriptomes). This may reflect non-detection of very divergent enzymes within this pathway, or equally that this pathway is not completely functional in the kareniacean plastid. We note that this pathway is also incomplete in stramenopile plastids (with the exception of diatoms; Nonoyama et al., 2019).

#### *S6.3: Fatty acid and plastid lipid biosynthesis*

Retention of a plastid-derived type II FAS pathway has already been implied in fucoxanthin and in peridinin dinoflagellates (Janouskovec et al., 2017) and our data supports this with all steps of this pathway identified in at least some of the species. Most of these enzymes are of plastid-late origin or have a plastid-late copy, but some additionally possess plastid-early homologues. The origins are not always consistent between the genera, for example FabI has a plastid-early copy in all except *T. helix* where the enzyme is of plastid-late origin while almost the opposite pattern can be observed for FabG.

The synthesis of sulfoquinovosyldiacylglycerol (SQDG) is the most striking part of the reconstructed kareniacean lipid metabolism as one of its enzymes, SQD2 is of green algal-like origin (Figure S4.3) in all genera and represents a very reliable case of conserved LGT, while the other enzyme SQD1 was not identifiable in any of the seven transcriptomes. SQDG is essential for plastid function and no previous lipidomic study suggests it is absent in kareniacean or other dinoflagellates (Leblond and

Chapman, 2000; Leblond et al., 2003), it was previously noted that the SQD1 (and in some cases also SQD2) is not detectable in the sequence data for *K. brevis*, *K. micrum*, and various other non-fucoxanthin dinoflagellates and other algae (Riccio et al., 2020), and in the proteome of isolated plastids of *Euglena* (Novák Vanclová et al., 2020). This may be due to constantly very low expression level of the gene to the point of not being captured during sequencing or existence of a yet undiscovered, non-homologous alternative enzyme.

#### *S6.4: Photosynthesis*

The majority of photosystem and light-harvesting complex subunits and parts of the photosynthetic electron transport chain are of plastid-late origin. PsaD is the only photosystem subunit that was vertically inherited from the ancestral plastid in *Karlodinium* and likely *Takayama* (where the protein was not predicted as plastid-targeted, likely due to N-terminal truncation) while it remains plastid-coded in *Karenia* (Dorrell et al., 2016). Another part of the photosynthetic apparatus that is not of plastid-late origin is plastidial ferredoxin (PetF) which shows a plastid-early origin in *Karenia* and *Karlodinium* and replaced a green-like homolog in *Takayama*. Of note, petF is also associated with non-photosynthetic metabolism (e.g., in leucoplasts of non-photosynthesizing plant tissues), and is plastid-encoded in the non-photosynthetic chrysophyte *Spumella* NIES-1486 (Dorrell et al., 2019), and its specific metabolic functions in Kareniaceae remain to be determined.

Most subunits of plastidial F-type ATP synthase are plastid-encoded in Kareniaceae and their sequences were not recovered in the transcriptomes. The gamma subunit (atpG, K02115) is encoded in the nucleus and is plastid-late origin in all species. Surprisingly, the delta subunit (atpH, K02113), an essential component of the complex which is plastid-encoded in most organisms but not in *K. micrum* and *K. mikimotoi*, was identified in neither the predicted plastid proteome nor the whole transcriptome of any of the studied organisms. The most conservative explanation for this could be that the gene is indeed plastid-encoded in kareniaceans (as it is in at least some haptophytes, Puerta et al., 2005) and was simply missed during plastid genome sequencing and assembly. However, a targeted HMMER search for possible distant homologs revealed that the distantly related functional analog of this protein in mitochondrial F-type ATP synthase (ATP5D, K02134) is duplicated in all species except *Takayama*, and in most of these extra copies, an N-terminal extension (in a few cases predictable as signal peptide, Figure S6.4.1) is present. This could indicate that a duplicated version of the mitochondrial protein was recruited and re-targeted to plastid to compensate for the loss of the analogous subunit of plastidial ATP synthase. This arrangement would be unprecedented as far as the current state of knowledge.

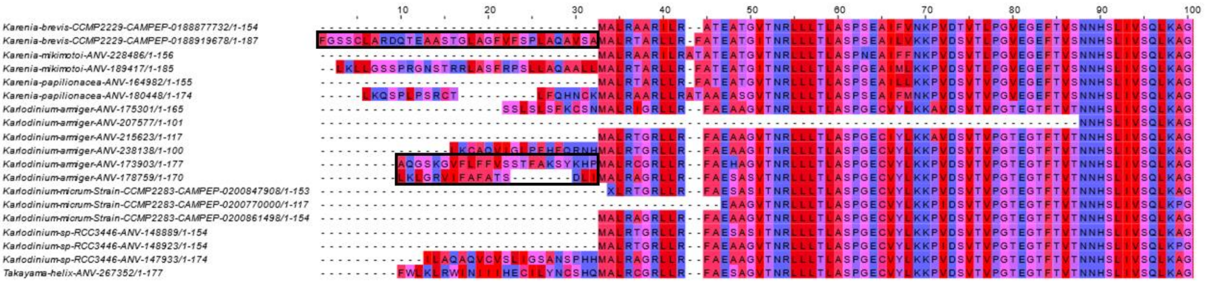


*Figure S6.4.1*: ClustalW alignment of the first 100 aa of the ATP5D sequences retrieved from the whole transcriptomes, coloured by hydrophobicity; N-terminal extension is present in at least one copy in all species except *Takayama*; in cases of *Karenia brevis* and *Karlodinium armiger* it is predictable as signal peptide (highlighted with black border).

*S6.5: Calvin cycle*

The Calvin cycle in kareniaceans is a true evolutionary mixture with most enzymes having multiple copies of both early and late origins. Sedoheptulose-bisphosphatase is the only protein that seems to be of purely plastid-late origin in all species, which may reflect that it is redox-regulated and has exclusive functions in photosynthesis. In contrast, triosephosphate isomerase and transketolase are exclusively of plastid-early origin, while further enzymes that function reversibly in both the glycolysis and pentose phosphate pathway possess both plastid-late and plastid-early copies. Notably, phosphoribulokinase is an exception from this overall trend as it is a photosynthetic carbon fixation-specific enzyme that possesses uniquely plastid-early origins.

*S6.6: Amino acid, nitrogen and sulfur metabolism*

While enzymes for the synthesis of most amino acids are present in the transcriptomes, only a small portion of them were predicted as plastid-targeted: the synthesis of alanine, aspartate, and asparagine from pyruvate and several following enzymes in the pathway towards lysine (up to 4-hydroxy-tetrahydrodipicolinate reductase), some of the enzymes converting glutamine to ornithine, the final step of proline synthesis (pyrroline-5-carboxylate reductase), some enzymes for serine and glycine synthesis from 3-phosphoglycerate, and the GS/GOGAT pathway. This low participation of plastid on amino acid synthesis is reminiscent of the situation in *Euglena* where only several isolated enzymes of serine and cysteine metabolism reside in the organelle (Novák Vanclová et al., 2020) as opposed to other algal groups, such as ochrophytes where the plastid is the site of synthesis of cysteine, aromatic, and branched-chain amino acids (Dorrell et al., 2017).

The shikimate pathway does not seem to be plastid-localized with the exception of the AROM polypeptide of *K. brevis* nand 3-dehydroquinate synthase which was predicted as plastidial in five of the species and exhibits a differential origin: plastid-early in *Karlodinium* and *Takayama* but plastid-late in *Karenia*. Interestingly, the same evolutionary division can be observed in the case for two of the enzymes of ornithine synthesis from glutamate, acetylglutamate kinase and N-acetyl-gamma-glutamyl-phosphate reductase. This may reflect the overall higher amount of plastid-late signal in *Karenia* in comparison to the other two genera.

Sulfur metabolism, including enzymes converting serine to cysteine, is evolutionarily mixed but comparatively richer in plastid-late signals. Most notably, the SUF system is not uniform in its origin and comprises of plastid-early SufS and SufB but plastid-late SufC, D, and E. Most of the nitrogen fixation enzymes were not predicted as plastid-targeted but the ones present are also evolutionarily non-uniform: while the putative plastidial nitrate and nitrite transporter and nitrate reductase are pan-alveolate (shared with ciliates) and plastid-early, respectively, nitrite reductase was gained with the new plastid.

*S6.7: Protein import and folding*

As expected, the protein import machinery was predominantly inherited from the current plastid donor as SELMA (the Symbiont-specific ERAD-Like MAchinery for protein import) is present in haptophytes and kareniaceans but not peridinin dinoflagellates, but there are a few notable exceptions. There are isolated cases of plastid-early SELMA components targeted to the plastid, although these may equally relate to host ERAD components with false positive predictions.

Alb3 represents an interesting case of a protein closely associated with plastid biogenesis and photosynthesis (since it plays a significant role in the insertion of photosynthetic complexes into the thylakoid membrane) with both plastid-early and plastid-late homologs in *K. brevis* and only a plastid-early homolog in *K. micrum* and *T. helix*.

*S6.8: Ribosome and aminoacyl-tRNA synthesis*

Nucleus-encoded and plastid-targeted ribosomal proteins were mostly gained with the current plastid with only a few exceptions. These include L7/L12 in *Karlodinium* and *Takayama*, L17 of *Karlodinium*, and L15 in *K.mikimotoi* and *K. armiger*, and might represent either genes retained from the ancestral plastid, or retargeted, originally mitochondrial proteins (e.g. L15 of *K. armiger* is shared not only with dinoflagellate but also apicomplexans). All three species of *Karenia* contain an additional copy of ribosomal protein S1 that seems to be of prokaryote origin (and shared with only a few eukaryotic algae) and was likely laterally gained. Single gene trees of these proteins indicate the presence of further, non-plastid targeted homologues in all *Karenia* transcriptomes, of as yet unclear function.

The plastidial inventory of aminoacyl-tRNA synthetases is evolutionarily mixed, with relatively high concentration of LGT. There are two potential cases of LGT from green algae, one in *Takayama* only (PARS, proline-tRNA synthetase) and a second (HARS, histidine-tRNA synthetase, Figure S5.1) in all except *Takayama* where, in turn, the protein seems to be brown in origin. Three other synthetases (TARS, tryptophanyl- (Figure S5.2), NARS, asparaginyl-, and MARS, methionyl-) are in turn of brown origin in at least some representatives.

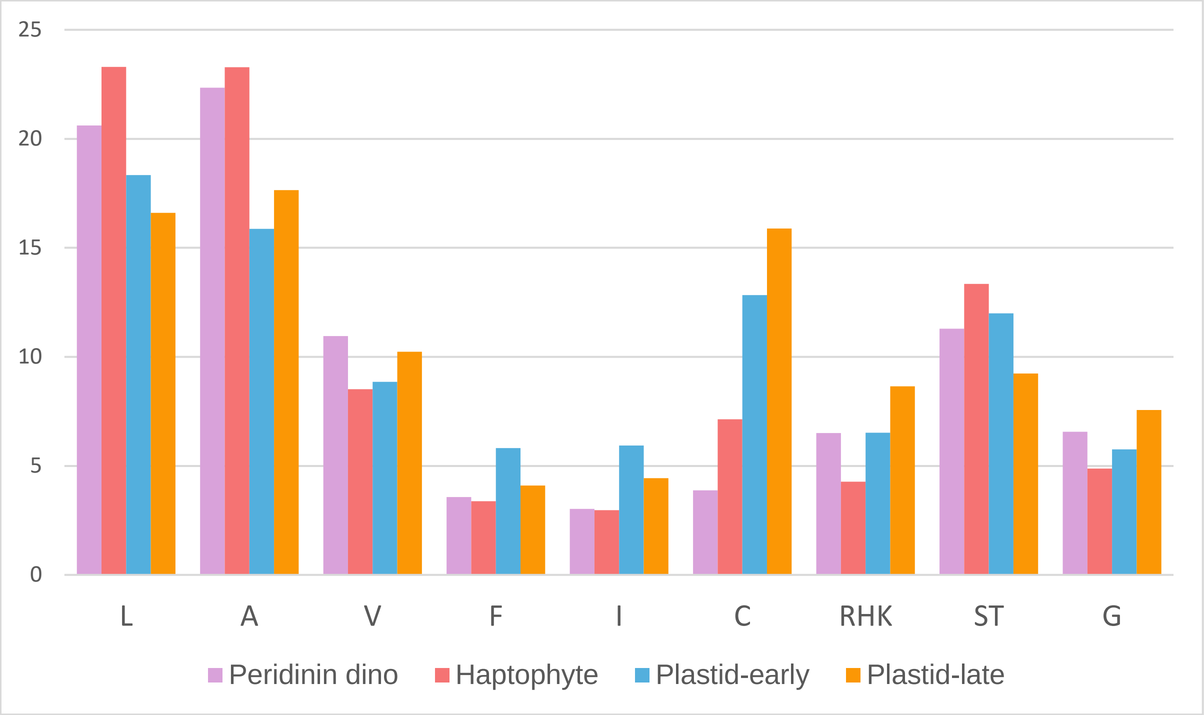


*Figure S7*: Amino acid composition of signal peptides of the putative plastidial proteins of five peridinin dinoflagellates, five haptophytes, and plastid-early and plastid-late proteins of the seven investigated kareniaceans. Hydrophobic residues dominate with A and L more slightly more frequent in peridinin dinoflagellates and haptophytes and F and I slightly more frequent in kareniaceans. The most notable is the enrichment of C in kareniaceans, especially in plastid-late proteins, reflective of the LACLAC motif unique to kareniaceans and especially conserved in their plastid-late proteins.

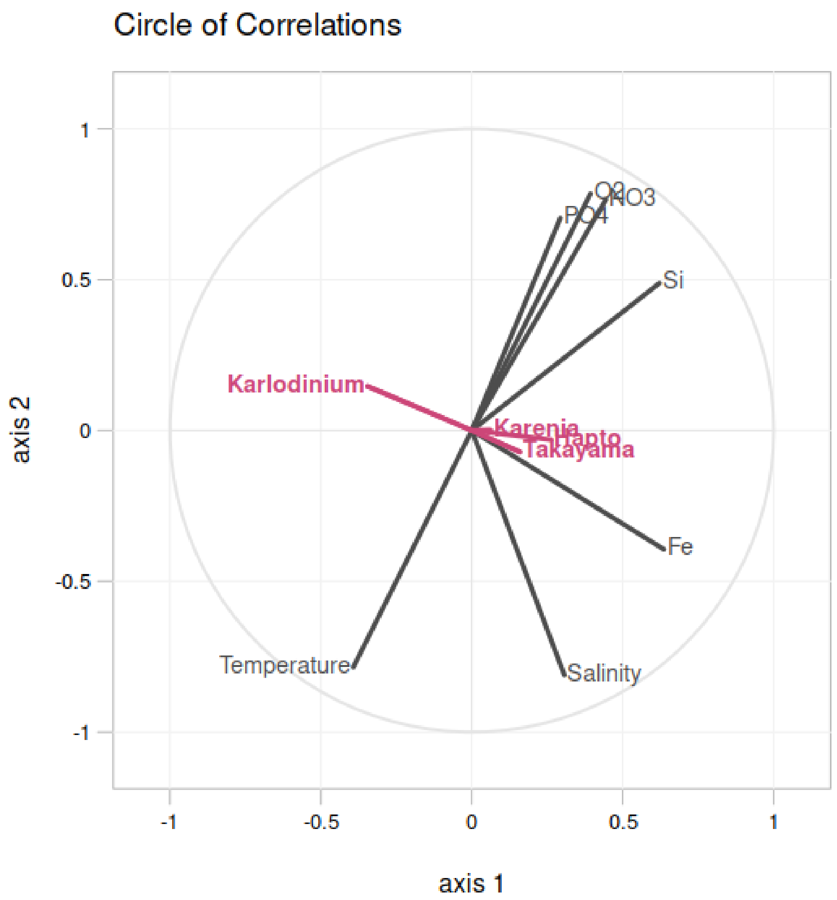


*Figure S8*: Partial Least Square analysis of environmental data and abundances of the three studied kareniacean genera and haptophytes in 75 *Tara* Oceans stations in which they occur (surface depth only): while haptophytes, *Karenia* and *Takayama* exhibit a similar correlation pattern (namely positive correlation with salinity, silica, and iron), *Karlodinium* shows the exact opposite trend.

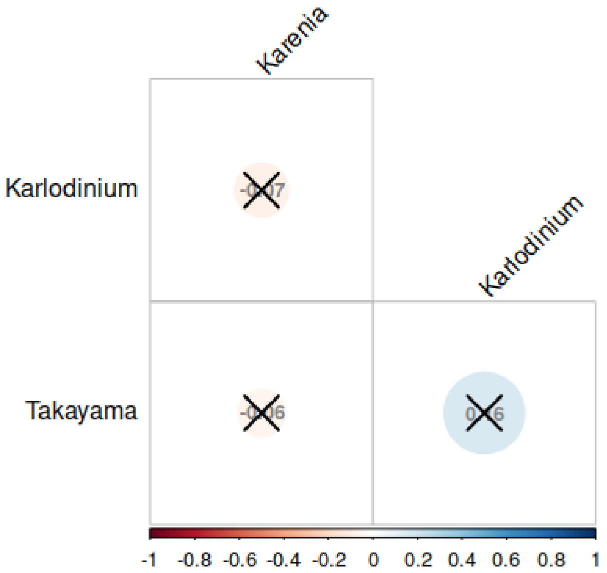


*Figure S9*: Correlation matrix (Spearman's rho correlation coefficient) of the three kareniacean genera abundances in 75 *Tara* Oceans stations in which at least one of the organisms was retrieved (pooled size fractions and depths): there is no significant correlation between either pair.

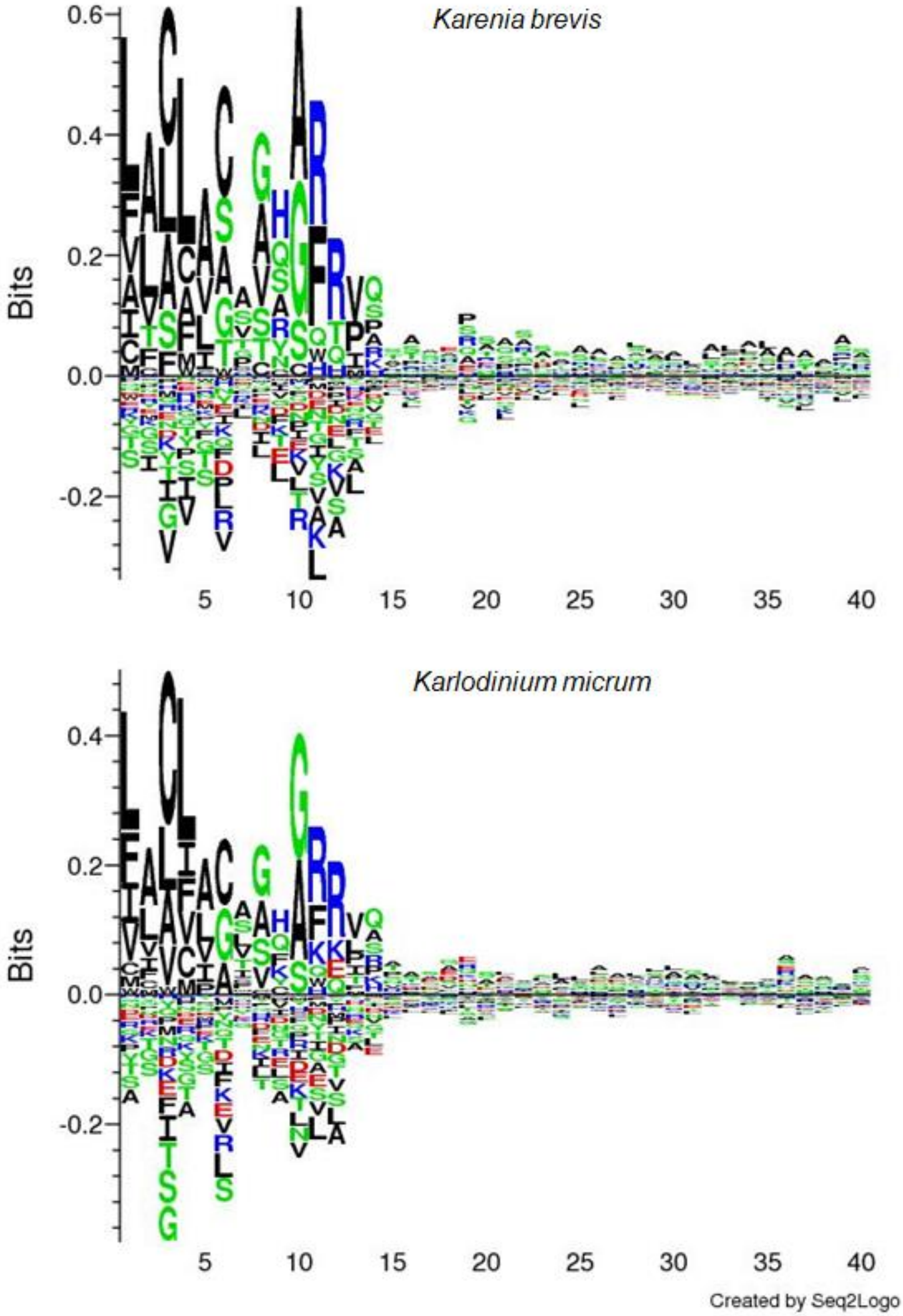


*Figure S10*: Sequence logos of the N-terminal region of the model plastidial datasets for *Karenia brevis* and *Karlodinium micrum* based on which the scoring matrix was prepared; the signal peptide cleavage position is between positions 10 and 11.

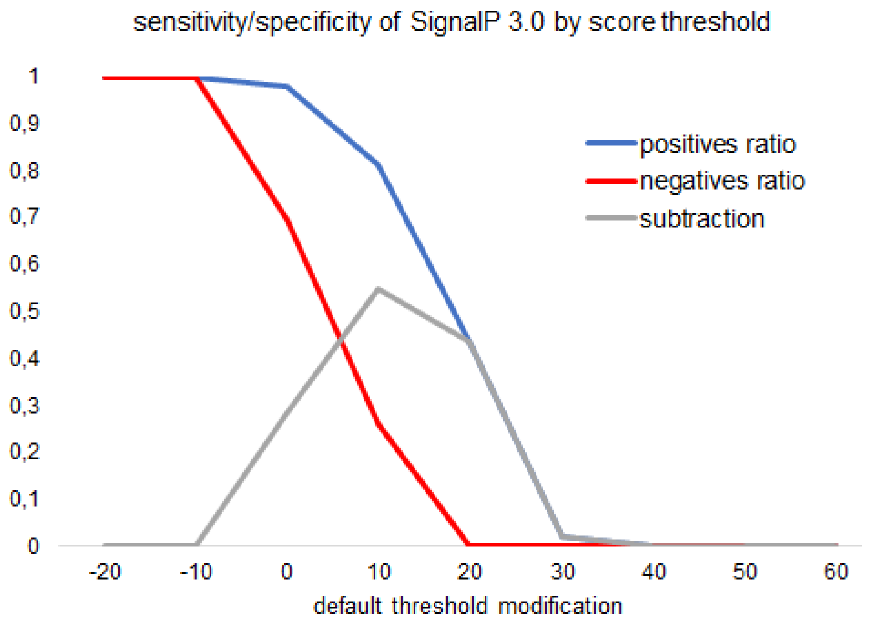


*Figure S11*: The effect of SignalP 3.0 score threshold modification on the specificity/sensitivity of prediction on model kareniacean datasets.

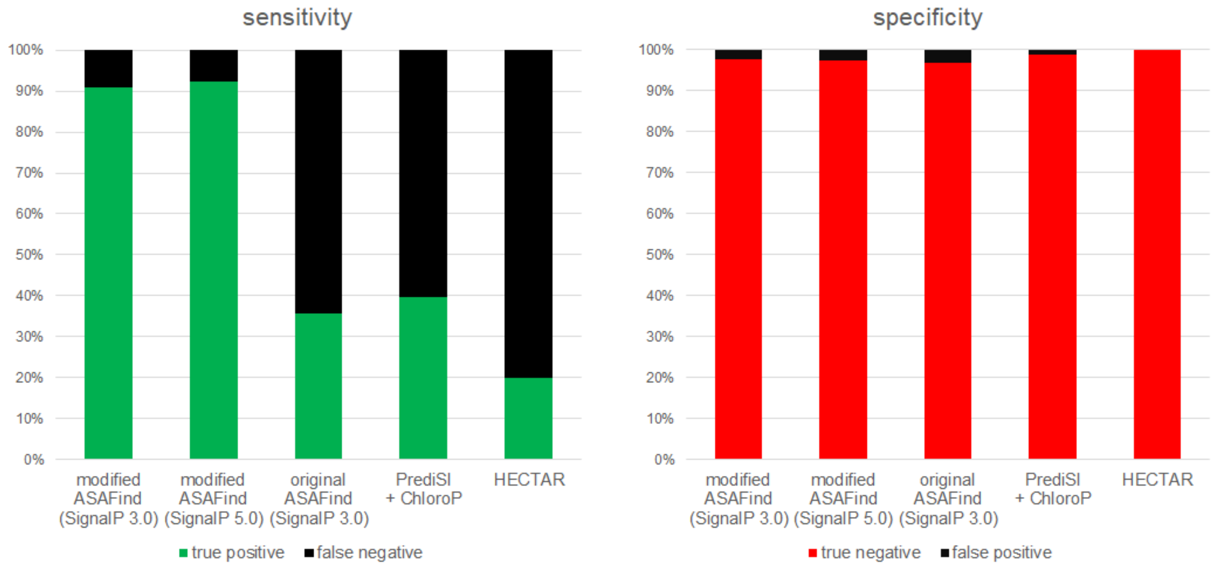


*Figure S12*: Sensitivity and specificity of the prediction software and their combinations tested during prediction optimization; while all methods exhibit comparable specificity, the sensitivity achieved by modified ASAFind script is much higher, particularly in conjunction with SignalP 5.0.
