## Supplementary material for "Divergent and diversified proteome content across a serially acquired plastid lineage": modASAFind: modASAFind.py

```
#! /usr/bin/python
#
# This work is copyright Cedar McKay and Gabrielle Rocap, University of Washington. This work
# is licensed under the Creative Commons Attribution-ShareAlike 3.0 Unported License. To view a
# copy of this license, visit http://creativecommons.org/licenses/by-sa/3.0/ or send a letter
# to Creative Commons, 171 Second Street, Suite 300, San Francisco, California, 94105, USA.
# 
# 
# ASAFind.py requires > Python 2.7 with Biopython. To install ASAFind, move this file to a
# directory in your PATH (such as /usr/local/bin), adjust the shebang path (first line) to your
# system's python path, and install Biopython. Developed using Biopython version 1.63.


import sys
import os.path
from collections import defaultdict
from optparse import OptionParser
from Bio import SeqIO

VERSION = "1.1.7"
PROJ = "ASAFind"

#### Collect Input ####
#######################

usage="""{} {}
%prog -f FILE -p FILE [options]

Takes a Fasta and companion SignalP (versions 3.0, 4.0 or 4.1) short format file as input,
either with the complete SignalP header (two lines starting with '#') or just with the results
(first line starts with first sequence name). SignalP truncates the sequence names to 20
characters. Therefore, ASAFind only considers the first 20 characters of the fasta name, which
must be unique within the file. Parts of the fasta name after the 20th character are ignored.
Additionally, the fasta name may not contain a '-' or '|'. This requirement is because SignalP
converts characters in sequence names (e.g. '-' is changed to '_'). ASAFind requires at least 7
aa upstream and 22 aa downstream of the cleavage site suggested by SignalP. The output of this
script is a tab delimited table.""".format(PROJ, VERSION)


parser = OptionParser(usage=usage, version="{} {}".format(PROJ, VERSION))


parser.add_option("-f", "--fasta_file", metavar="FILE", dest="fasta_file", default=None,
                help="Specify the input fasta FILE.")

parser.add_option("-p", "--signalp_file", metavar="FILE", dest="signalp_file", default=None,
                help="Specify the input SignalP FILE.")

parser.add_option("-t", "--score_table_file", metavar="FILE", dest="score_table_file",
                default=None, help="Optionally, specify a custom scoring table.")

parser.add_option("-s", "--short", metavar="FILE", dest="short_output",
                default=False, action='store_true', help="Short format output")
    
parser.add_option("-o", "--out_file", metavar="FILE", dest="out_file", default=None,
                help="Specify the path and name of the output fasta file you wish to create."
                "Default will be the same as the fasta_file, but with a '.tab' suffix.")

parser.add_option("-w", "--web_output", dest="web_output",
                default=False, action='store_true', help="Format output for web display. This is "
                        "mostly useful when called by a web app")


(options, args) = parser.parse_args()


#### Variables and Names ####
#############################

#Figure out some names and paths
if options.fasta_file and options.signalp_file:
    fasta_file = os.path.abspath(options.fasta_file)
    signalp_file = os.path.abspath(options.signalp_file)
else:   
    print("{0} {1}. Type '{0}.py -h' for help.".format(PROJ, VERSION))
    sys.exit()

(fasta_file_path, fasta_file_name) = os.path.split(fasta_file)
(fasta_file_base_name, fasta_file_ext) = os.path.splitext(fasta_file_name)


if options.score_table_file:
    custom_scoring_table = open(options.score_table_file, 'rU').read()
else:
    custom_scoring_table = None

short_output = options.short_output
web_output = options.web_output

#Figure out what our out_file is.
if options.out_file:
    out_file = os.path.abspath(options.out_file)
else:
    out_file = os.path.abspath(os.path.join(fasta_file_path, fasta_file_base_name + '.tab'))


#### Functions ####
###################

def parse_signalP(p_file):
    '''Processes a signalP output file, and returns a parsed dictionary of results
    
    Args:
        Path to a signalP 3.0 format file, short format, both eukNN and HMM
    
    Returns:
         Dict with the keys of Cmax, Ymax, Smax, Smean, D. Each value is a tuple
         (score, signalp-Y/N, position). Smean and D don't have a position.
    
    '''
    # name Cmax pos ? Ymax pos ? Smax pos ? Smean ?  D  ?  name !  Cmax pos ? Sprob ?
    #  0    1    2  3  4    5  6  7    8  9  10  11 12  13  14  15  16   17 18  19  20
    d = {} #results dict
    p_handle = open(p_file, 'rU')
    seen = set()
    
    
    #Account for different column order. Thanks guys. 
    line = next(p_handle) #First line
    if line.startswith('# SignalP-NN') or len(line.split()) == 21:
        d['signalp_version'] = 'SignalP-3.0'
        Cmax_score = 1
        Cmax_position = 2
        Ymax_score = 4
        Ymax_position = 5
        Smax_score = 7
        Smax_position = 8
        Smean_score = 10
        D_score = 12
        D_conclusion = 13
        
    elif line.startswith('# SignalP-4') or len(line.split()) == 12:
        if line.startswith('# SignalP-4'):
            d['signalp_version'] = line.split()[1] #Right now this can be 4.0 or 4.1
        else:
            d['signalp_version'] = 'SignalP-4.x'
        Cmax_score = 1
        Cmax_position = 2
        Ymax_score = 3
        Ymax_position = 4
        Smax_score = 5
        Smax_position = 6
        Smean_score = 7
        D_score = 8
        D_conclusion = 9
    else:
        raise Exception('The SignalP file is an unrecognized format')


    p_handle.seek(0) #In case we didn't have a header, we need to go back to beginning of file.
    for line in p_handle:
        if not line.startswith("#") and len(line) > 1:
            atoms = line.split()
            id = atoms[0][:20]
            if id in seen:
                raise Exception('The SignalP file has entries not unique within first 20 '
                                'characters: {}'.format(id))
            else:
                seen.add(id)
            d[id] = {'Cmax':{'score': atoms[Cmax_score],'position':
            int(atoms[Cmax_position])}, 'Ymax':{'score': atoms[Ymax_score],
            'position':int(atoms[Ymax_position])}, 'Smax':{'score': atoms[Smax_score],
            'position':int(atoms[Smax_position])}, 'Smean':{'score': atoms[Smean_score]},
            'D':{'score': float(atoms[D_score]), 'conclusion': atoms[D_conclusion]}}
                                        
    
    return d


def parse_score_table(scoring_table_string = None):
    '''Takes a scoring table, and returns a 2D dict (like a lookup table)
    
    The input table should be in the following format:
    
    A   0.151009115 0.085083382 0.583560009
    C   0.031243265 0.026179502 0.063661092
    D   0.005207211 0   0

    For example, the score weight given to a 'C' in the second position is 0.026179502
    
    Args:
        Scoring table as a string. If none, the default table will be used.
    
    Returns:
        A 2D dict describing the table. Access the scoring weight to a 'C' in the second
        position like this: d['C'][2]. Note position is not zero based, to match SignalP
        results.
    '''

    if not scoring_table_string:
        scoring_table_string = \
            '''A    0.151009115 0.085083382 0.583560009 0.090317708 2.280392482 0   0.347218528 0.059318647 0.097200366 0.074468452 0.033263496 0.041608564 0.056157785 0.095102845 0.071684231 0.043794002 0.079695775 0.085701177 0.044777601 0.057920154 0.059972166 0.114027446 0.075960804 0.061762271 0.047797224
            C   0.031243265 0.026179502 0.063661092 0.009507127 0.074159105 0   0.017360926 0.009886441 0.004860018 0.004380497 0.003695944 0   0   0   0.010240604 0   0.004980986 0.005356324 0   0   0   0.004072409 0   0.003250646 0.00896198
            D   0.005207211 0   0   0.028521381 0   0   0   0   0.009720037 0.004380497 0   0   0   0   0.005120302 0.0043794   0   0.010712647 0.008141382 0.010860029 0.014993042 0.012217226 0.011394121 0.013002583 0.023898612
            E   0.015621633 0.013089751 0   0.06654989  0   0   0.008680463 0   0   0.004380497 0   0.004623174 0.003509862 0   0.005120302 0   0   0   0   0   0.011244781 0.020362044 0.015192161 0.019503875 0.023898612
            F   0.015621633 0.006544876 0   0.014260691 0   3.251039844 0   0.059318647 0.009720037 0.017521989 0.059135103 0.023115869 0.031588754 0.02237714  0.005120302 0.0131382   0.029885915 0.032137942 0.016282764 0.014480038 0.011244781 0.012217226 0.015192161 0.016253229 0.011949306
            G   0.104144217 0.065448755 0.05305091  0.019014254 0.333715973 0.021819059 0.02604139  0.049432206 0.024300091 0.03066348  0.022175664 0.069347606 0.049138061 0.02237714  0.025601511 0.017517601 0.039847887 0.016068971 0.012212073 0.021720058 0.011244781 0.020362044 0.034182362 0.029255813 0.014936633
            H   0.026036054 0.019634627 0   0.057042763 0   0   0.02604139  0.029659324 0.02916011  0.03066348  0.022175664 0.009246348 0.010529585 0.02237714  0.015360907 0.026276401 0.029885915 0.016068971 0.028494837 0.021720058 0.018741302 0.004072409 0.011394121 0.003250646 0.005974653
            I   0   0.019634627 0.095491638 0.009507127 0   0.021819059 0.069443706 0   0.019440073 0.008760994 0.01847972  0.023115869 0.017549308 0.039159995 0.010240604 0.017517601 0.029885915 0.021425294 0.020353455 0.025340067 0.011244781 0.016289635 0   0.006501292 0.005974653
            K   0.015621633 0.006544876 0   0   0   0   0.008680463 0   0.004860018 0.021902486 0.007391888 0.013869521 0.028078892 0.02237714  0.025601511 0.017517601 0.029885915 0.016068971 0.012212073 0.018100048 0.018741302 0.012217226 0.026586281 0.013002583 0.023898612
            L   0.031243265 0.078538506 0.021220364 0.047535636 0   0.065457178 0.008680463 0.138410177 0.048600183 0.03066348  0.070222935 0.046231738 0.031588754 0.039159995 0.051203022 0.061311602 0.039847887 0.117839119 0.093625892 0.061540163 0.063720426 0.03257927  0.053172563 0.052010334 0.023898612
            M   0.010414422 0.013089751 0   0.004753564 0   0   0.017360926 0   0.019440073 0.013141491 0.007391888 0   0.007019723 0.016782855 0   0.0087588   0   0   0.008141382 0.00362001  0.018741302 0.012217226 0.00759608  0.013002583 0.014936633
            N   0.015621633 0.026179502 0.010610182 0.071303454 0   0.021819059 0.008680463 0.029659324 0.02916011  0.021902486 0.03695944  0.03698539  0.010529585 0.01118857  0.020481209 0.017517601 0.009961972 0.010712647 0.028494837 0.036200096 0.022489562 0.024434453 0.041778442 0.019503875 0.026885939
            P   0.041657687 0.045814129 0   0   0.018539776 0   0.008680463 0.781028858 0.136080512 0.078848949 0.066526991 0.07397078  0.056157785 0.04475428  0.061443627 0.035035201 0.044828873 0.085701177 0.04070691  0.039820106 0.029986083 0.044796497 0.041778442 0.032506459 0.026885939
            Q   0.036450476 0.039269253 0   0.038028509 0   0   0.086804632 0.049432206 0.034020128 0.043804972 0.025871608 0.018492695 0.031588754 0.03356571  0.030721813 0.056932202 0.029885915 0.032137942 0.024424146 0.018100048 0.022489562 0.024434453 0.026586281 0.026005167 0.026885939
            R   0.005207211 0.006544876 0   0.014260691 0   0   0   0.009886441 0.034020128 0.03066348  0.051743216 0.064724433 0.052647923 0.139857124 0.128007556 0.100726203 0.089657746 0.058919559 0.044777601 0.047060125 0.037482604 0.044796497 0.022788241 0.026005167 0.014936633
            S   0.182252381 0.340333526 0.275864732 0.209156797 0.352255749 0   0.234372506 0.168069501 0.160380604 0.183980881 0.103486431 0.161811081 0.0912564   0.240554254 0.15872937  0.144520205 0.174334507 0.235678238 0.12619142  0.115840307 0.116196072 0.138461899 0.136729447 0.100770022 0.104556428
            T   0.114558639 0.222525767 0.360746187 0.061796326 0.018539776 0   0.329857601 0.177955942 0.087480329 0.087609943 0.051743216 0.11095617  0.052647923 0.111885699 0.143368463 0.100726203 0.144448591 0.085701177 0.089555201 0.065160173 0.078713468 0.069230949 0.056970603 0.065012917 0.05974653
            V   0.05727932  0.052359004 0.286474914 0.019014254 0   0   0.243052969 0.069205089 0.05832022  0.026282983 0.025871608 0.055478085 0.0456282   0.061537135 0.076804534 0.048173402 0.044828873 0.042850589 0.065131055 0.028960077 0.071216947 0.061086132 0.041778442 0.029255813 0.023898612
            W   0.005207211 0   0   0.004753564 0   0.152733416 0   0.009886441 0   0   0.003695944 0.009246348 0   0.005594285 0.005120302 0.0087588   0   0.016068971 0.004070691 0   0   0   0.00379804  0   0.005974653
            Y   0   0.019634627 0.010610182 0.023767818 0   0.087276237 0   0   0   0.013141491 0.003695944 0.004623174 0.007019723 0   0   0.0043794   0.004980986 0   0.008141382 0.014480038 0.00374826  0.008144818 0.00759608  0.009751938 0
            X   0   0   0   0   0   0   0   0   0   0   0   0   0   0   0   0   0   0   0   0   0   0   0   0   0
            *   0   0   0   0   0   0   0   0   0   0   0   0   0   0   0   0   0   0   0   0   0   0   0   0   0'''


    d = defaultdict(dict) #results dict
    for line in scoring_table_string.split('\n'):
        atoms = line.split()
        aa = atoms.pop(0) #amino acid short code
        d
        i = 1 #positions are 1 based
        while len(atoms) > 0:

            d[aa][i] = float(atoms.pop(0))
            i += 1
    
    return d
            
        
    

    


def score_peptide(seq, table = None):
    '''Takes a 25mer and calculates the cleavage site score and transit peptide score.
    
    Args:
        A 25 aa long string, and a 2D dict representation of a scoring table.

    Returns:
        A tuple of scores. First is cleavage site score, 2nd is transit peptide score. 
    '''
    
    if not table:
        raise Exception("Expected a scoring table as a 2D dict")
    if len(seq) != 25:
        raise Exception("Expected a 25mer")
    seq = seq.upper()
    cleavage_score = 0
    transit_score = 0

    i = 1 # 1 based counting conforming to signalP
    for aa in seq:
        cleavage_score += table[aa][i]
        
        if i > 9: #transit score is calculated omitting 1st 9 aa.
            transit_score += table[aa][i]
        i += 1
    
    return (cleavage_score, transit_score)


#### Main ####
##############

if sys.version_info < (2,7):
    print ("WARNING: This version of ASAFind was developed on Python 2.7. " \
          "It is not tested on Python {}.{}".format(sys.version_info[0], sys.version_info[1]))


if web_output:
    print("<PRE>")
    
#Test input files to make sure they are parseable, valid, no dups.
try:
    records_dict = SeqIO.to_dict(SeqIO.parse(fasta_file, "fasta"))
except ValueError as e:
    print("The input fasta could not be parsed. The error is: {}".format(e))
    raise e

    
try:
    signalps = parse_signalP(signalp_file) #It parses. That means no duplicate entries.

    records_keys_short = [x[:20] for x in list(records_dict.keys())]
    problem_names = [x for x in records_keys_short if any([s in x for s in ('-','|')])]
    if problem_names:
        print("Problem names: ", problem_names)
        raise Exception("SignalP changes '-' and '|' characters in names to '_', therefore ASAFind "
                         "does not allow those characters either.")

    if not len(records_keys_short) == len(set(records_keys_short)):
        dups = tuple(set([x for x in records_keys_short if records_keys_short.count(x) > 1]))
        raise Exception('The input fasta has records that are not unique to the first 20 '\
                        'characters. The offending records begin: {}'.format(dups))
    
    if not set(signalps.keys()) >= set(records_keys_short): 
        raise Exception('SignalP file does not have an entry (unique to first 20 '
                        'characters) for every sequence in fasta file. Missing entry for '
                        '{}'.format([x for x in set(records_keys_short) - set(signalps.keys())]))
except Exception as e:
    print('The SignalP file could not be validated. The error is:\n{}'.format(e))
    sys.exit()


#Input files passed our tests. Onward!
records = SeqIO.parse(fasta_file, "fasta")
signalps = parse_signalP(signalp_file) #This is a dict
scoring_table = parse_score_table(custom_scoring_table)

#For stats
total_proteins = 0
signalp_positive = 0
chloroplast_targeted = 0
chloroplast_targeted_high = 0
chloroplast_targeted_low = 0
skipped_proteins = 0

results = []
for record in records:
    total_proteins += 1
    signalp = signalps[record.name[0:20]] #Get a single parsed signalp result for the
                                          #record. SignalP only uses 20 chars of the name

    best_window = {'window': 0, 'score_25mer': -100, 'score_20mer': -100,
                   'peptide_25mer': 0}

    if signalp['D']['score'] > 0.35:
        signalp_positive += 1        
        if len(record.seq[signalp['Ymax']['position']-13 : signalp['Ymax']['position']+16]) < 29   :
            results.append((record.name,
                        'Y',
                        signalp['D']['score'],
                        'NA',
                        'NA',
                        'NA',
                        'NA',
                        'NA',
                        'NA',
                        'NA',
                        'NA',
                        'NA',
                        'No prediction, sequence too short',
                        str(record.seq)))
            
            skipped_proteins += 1
            continue
            
        sliding_window_range = (-2,-1,0,1,2) #Hardcode since no plans to change

        for window in sliding_window_range:
            #subtract 1 to convert to 0 based positions
            #25mer starts 5 before cleavage position.
            start = signalp['Ymax']['position'] + window - 10 - 1
            stop = start + 25
            peptide_25mer = record.seq[start:stop]


    
            # Score returned is a tuple. 0 is 25mer score and 1 is 20mer score
            this_window_score = score_peptide(peptide_25mer, scoring_table)
                                                          
            if this_window_score[0] > best_window['score_25mer']:
                best_window = {'window': window, 'score_25mer': this_window_score[0],
                               'score_20mer': this_window_score[1],
                               'peptide_25mer': peptide_25mer}
            
            if window == 0: #Store the original predicted window when we run across it.
                window_0 = {'window': window, 'score_25mer': this_window_score[0],
                            'score_20mer': this_window_score[1],
                            'peptide_25mer': peptide_25mer}
            
        #Check if 1st of cleaved peptide of best window starts with F,W,Y, L
        best_window['CSS'] = 0
        plus1_aa = best_window['peptide_25mer'][10].upper()
        if plus1_aa in 'RF':
            best_window['plus1_aa'] = plus1_aa
            best_window['CSS'] += 2
        else:
            best_window['plus1_aa'] = 'No'
        plus2_aa = best_window['peptide_25mer'][11].upper()
        if plus2_aa in 'RV':
            best_window['plus2_aa'] = plus2_aa
            best_window['CSS'] += 1
        else:
            best_window['plus2_aa'] = 'No'    
        plus3_aa = best_window['peptide_25mer'][2].upper()
        if plus3_aa in 'CALS':
            best_window['plus3_aa'] = plus3_aa
            best_window['CSS'] += 2
        else:
            best_window['plus3_aa'] = 'No'
        plus4_aa = best_window['peptide_25mer'][5].upper()
        if plus4_aa in 'CGS':
            best_window['plus4_aa'] = plus4_aa
            best_window['CSS'] += 1
        else:
            best_window['plus4_aa'] = 'No'  
        if best_window['CSS'] > 1:
          if best_window['window'] == 0 and best_window['score_20mer'] > 10: 
                best_window['prediction'] = 'Plastid, high confidence'
                chloroplast_targeted += 1
                chloroplast_targeted_high += 1
          else:
                best_window['prediction'] = 'Plastid, low confidence'
                chloroplast_targeted += 1
                chloroplast_targeted_low += 1
        else:
            best_window['prediction'] = 'Not plastid, SignalP positive'
       #Calculate a few values that would be too cumbersome to put right into the results list
        window_0['position'] = signalp['Ymax']['position']
        best_window['position'] = signalp['Ymax']['position'] + best_window['window']

        results.append((record.name,
                        'Y',
                        signalp['D']['score'],
                        window_0['position'],
                        window_0['score_25mer'],
                        window_0['peptide_25mer'],
                        best_window['position'],
                        best_window['score_25mer'],
                        best_window['peptide_25mer'],                   
                        best_window['plus1_aa'],
                        best_window['plus2_aa'],
                        best_window['plus3_aa'],
                        best_window['plus4_aa'],
                        best_window['CSS'],
                        best_window['window'],
                        best_window['score_20mer'],
                        best_window['prediction'],
                        str(record.seq)))
                    
    elif signalp['D']['conclusion'] == 'N':
        predicted_25mer = record.seq[signalp['Ymax']['position']: signalp['Ymax']['position'] + 25]
        results.append((record.name,
                        'N',
                        signalp['D']['score'],
                        'NA',
                        'NA',
                        'NA',
                        'NA',
                        'NA',
                        'NA',
                        'NA',
                        'NA',
                        'NA',
                        'Not plastid, SignalP negative',
                        str(record.seq)))
    
    else:
        raise Exception("Expected either a Y or N.")


headers =   ('Identifier',
            'SignalP', # yes or no
            'SignalP D score',
            'SignalP cleavage position',
            'SignalP 25aa cleavage score',
            'SignalP 25aa sequence',
            'ASAFind cleavage position',
            'ASAFind 25aa cleavage score',
            'ASAFind 25aa sequence',
            'ASAFind cleavage has [RF] at 10 position',
            'ASAFind cleavage has [RV] at 11 position',
            'ASAFind cleavage has [CALS] at 2 position',
            'ASAFind cleavage has [CGS] at 5 position',
            'Critical site score',
            'ASAFind/SignalP cleavage site offset',
            'ASAFind 20aa transit score',
            'ASAFind Prediction',
            'Protein sequence')
            

out_handle = open(out_file, 'w')

if short_output:
    headers = [headers[i] for i in (0, 1, 6, 10, 11, 12)]

out_handle.write('\t'.join(headers) + '\n')

for result in results:
    if short_output:
        result = [result[i] for i in (0, 1, 6, 10, 11, 12)]
    #convert every item in list to string, and round floats to 9 decimal places. 
    result = [str('{:.9f}'.format(x)) if type(x) is float else str(x) for x in result]
    out_handle.write('\t'.join(result) + '\n')
out_handle.close()

chloroplast_not_targeted = signalp_positive - chloroplast_targeted - skipped_proteins

print("""\nThis is ASAFind version {}.
You used {}.
You submitted {} proteins
{} of your proteins were SignalP negative
{} of your proteins were SignalP positive
    {} of these were predicted to go to the plastid
        {} of these were predicted with high confidence
        {} of these were predicted with with low confidence
    {} of these were predicted not to go to the plastid """.format(VERSION,
            signalps['signalp_version'], total_proteins, total_proteins-signalp_positive,
            signalp_positive, chloroplast_targeted, chloroplast_targeted_high,
            chloroplast_targeted_low, chloroplast_not_targeted))
    
    
if skipped_proteins:
    print("    {} could not be analyzed because the length is too short".format(skipped_proteins))

    
print ("""\nCitation: If you use ASAFind in your research please cite our publication (Gruber et al., doi:
10.1111/tpj.12734) as well as the publications for the appropriate SignalP versions (Petersen
et al., doi: 10.1038/nmeth.1701 for SignalP 4.1 or Bendtsen et al., doi:
10.1016/j.jmb.2004.05.028 for SignalP 3.0).
""")

if web_output:
    print("</PRE>")
else:
    print ("Wrote results to {}".format(out_file))
```
