## Supplementary material for "Divergent and diversified proteome content across a serially acquired plastid lineage": SP5toSP4: SP5toSP4.py

```
#!/bin/bash/python2.7

import sys
import getopt

args = sys.argv[1:]

infilename = "input.txt"
outfilename = "output.txt"

if len(args) < 1:
    pass
else:
    infilename = args[0]
    
if len(args) < 2:
    pass
else:
    outfilename = args[1]

out = open(outfilename, "w")

def find_nth(line, char, n):
    start = line.find(char)
    while start >= 0 and n > 1:
        start = line.find(char, start+1)
        n -= 1
    return start

out.write("# SignalP-4.1 euk predictions\n# name\tCmax\tpos\tYmax\tpos\tSmax\tpos\tSmean\tD\t?\tDmaxcut\tNetworks-used\n")

with open(infilename) as file:
	for line in file:
		if line.startswith("#"):
			pass
		else:
			name = line[:line.find("\t")]
			score = line[find_nth(line, "\t", 2)+1:find_nth(line, "\t", 3)]
			if "SP(Sec/SPI)" in line:
				decision = "Y"
				position = line[line.find("CS pos: ")+8:line.find("-")]
			else:
				position = "0"
				decision = "N"
			newline = name + "\t" + score + "\t" + position + "\t" + score + "\t" + position + "\t" + score + "\t" + position + "\t" + score + "\t" + score + "\t" + decision + "\t0.450\tSignalp-noTM\n"
			out.write(newline)
		name = ""
		score = ""
		positin = ""
		decision = ""
		newline = ""
out.close()
```
